# Adipose Tissue Inflammation, Oxidative Stress, and Altered Adipogenesis Are Drivers of Dyslipidemia – A Multi-Omics Overview of Dyslipidemia in Obesity

**DOI:** 10.1101/2025.10.15.682716

**Authors:** Max S.Z. Zwartjes, Patrick A. de Jonge, Arnold W. van de Laar, Sjoerd C. Bruin, Abraham S. Meijnikman, Albert K. Groen, Victor E.A. Gerdes, Max Nieuwdorp

## Abstract

**Background:** Obesity is an important risk factor for cardiometabolic disease including dyslipidemia and atherosclerotic cardiovascular disease. While the role of the liver in dyslipidemia is established, the contribution of adipose tissue is less clear. This study aims to clarify the role of adipose tissue in lipid metabolism and its impact on dyslipidemia development.

**Methods:** We conducted a cross-sectional analysis of 125 patients from the BARIA longitudinal cohort study, consisting of patients with obesity undergoing bariatric surgery. Comprehensive phenotyping with fasting untargeted plasma metabolomics and lipid, lipoprotein, adipokine, RNA sequencing, and fecal sample shotgun metagenomic analysis was performed. We then compared tissue transcriptomic and plasma metabolites in individuals with and without dyslipidemia, exploring protective and pathogenic mechanisms.

**Results:** Dyslipidemia was present in 43 patients (34.4%), with elevations in triglycerides, ApoB, and reductions in HDL-C and ApoAI. Plasma adipokines were less reliable markers of dyslipidemia: leptin levels were unexpectedly reduced in dyslipidemia, and after adjusting for gender, age, and body weight, adipokines did not differ between patients with and without dyslipidemia. In relation to plasma lipids or dyslipidemia, RNA sequencing identified altered gene expression of liver, jejunum, visceral and subcutaneous adipose tissue. Adipose tissue of patients with dyslipidemia was characterized by gene alterations of 3 pathways: inflammation, oxidative stress, and adipogenesis. Untargeted plasma metabolomics revealed associations of plasma lipids with endocannabinoid-like, secondary bile acid, plasmalogen, butyrate, and sphingolipid metabolites. Gut metagenome analysis in dyslipidemia found only correlation on bacterial order level.

**Conclusions:** This study provides novel insights into the role of adipose tissue in dyslipidemia of obesity. Our findings highlight the importance of not only visceral, but also subcutaneous adipose tissue in regulating plasma lipids. We found that fasting dyslipidemia associated with adipose tissue inflammation, oxidative stress, and altered adipogenesis, providing new insights into metabolic regulation of plasma lipids beyond hepatic pathways.

## Introduction

Obesity is increasing worldwide and for individuals with a high BMI, cardiovascular disease remains the most attributed cause of death.^1,2^ Adipose tissue (AT) is not only an energy storing organ, but also a metabolic and endocrinologically active organ and a driver of cardiometabolic diseases like insulin resistance and type 2 diabetes (T2D), hypertension, metabolic dysfunction-associated steatotic liver disease (MASLD), and atherosclerotic cardiovascular disease (ASCVD).^3–5^ Obesity not only contributes to insulin resistance, but also directly to dyslipidemia, marked by decreased HDL-C and increased plasma TG concentration through enhanced hepatic release of apolipoprotein B-containing lipoproteins, such as VLDL.^6^ The mechanism behind the development of dyslipidemia in obesity is only partly understood, and studies have shown that degree of obesity and insulin resistance does not have a linear relationship with dyslipidemia; individuals with overweight (BMI 25.0-29.9 kg/m^2^) may have more significant changes in their lipid profile as compared to individuals with class III obesity (BMI over 40 kg/m^2^).^7,8^ This indicates that besides pathogenic mechanisms, also protective ones could be at play in obesity, influencing susceptibility to develop dyslipidemia and subsequent ASCVD.

While the role of the liver in lipid metabolism is undisputed, AT is increasingly seen as another important player.^9^ According to population-based and cohort studies, the amount of visceral adipose tissue (VAT) appears to be a stronger factor for the development of dyslipidemia and associated CVD risk than subcutaneous adipose tissue (SAT).^10–12^ Apart from the volume of AT, the function of AT, or rather, AT *dysfunction*, appears to influence glucose and lipid metabolism most.^4^ AT *dysfunction* is marked not only by adipocyte hypertrophy, but also by AT inflammation (M1-macrophage predominance), oxidative stress, and alterations in the adipokine profile such as increased leptin and decreased adiponectin concentrations.^6,13–15^ Changes in adipokine release is relevant, because, on the one hand, adipokines such as adiponectin, omentin and FGF-21 have insulin sensitizing, anti-inflammatory, and anti-atherogenic properties, whereas leptin, resistin and visfatin are thought to be detrimental for health.^9,14,16^ One of the suggested mechanisms contributing to a dysfunctional AT is hypoxia and oxidative stress caused by AT hypertrophy exceeding its vascularization potential.^17^

As mentioned, not all individuals with excess of AT develop dyslipidemia, hence further characterization of the role of AT in lipid metabolism is necessary. By comparing AT and liver gene expression patterns and metabolites of individuals with obesity and dyslipidemia versus individuals with obesity but without dyslipidemia, both protective and pathogenic mechanisms by which AT interacts with other tissues can be uncovered. Gut microbiota and its produced metabolites are also potential players in obesity, dyslipidemia, and ASCVD.^6,18^

Therefore, in this cross-sectional study we hypothesized that besides liver, also different AT depots (specifically VAT), have significant up- or downregulated genetic pathways in association with dyslipidemia. We thus postulate that AT in individuals with obesity can undergo one of two distinct pathways: one involving excess of AT (hyperplasia) but with maintenance of lipid homeostasis, while another involves AT dysfunction and the development of dyslipidemia. We studied this hypothesis in a cohort of morbidly obese patients scheduled for bariatric surgery in which comprehensive metabolic phenotyping was performed, including measurements of insulin resistance, tissue RNA, plasma metabolites, lipids, and adipokines, as well as determination of the gut microbiota composition. We aimed to gain insight into pathways and genes involved in dyslipidemia in individuals with obesity, specifically those of VAT and SAT, and we specifically focused on genes and adipokines proposed to be involved in AT dysfunction such as inflammation and oxidative stress.

## Methods

### Study design and population

We used blood-, fecal, and tissue biopsies of jejunum, liver, as well as adipose tissue (subcutaneous and visceral) obtained during surgery from selected patients of the BARIA longitudinal cohort study.^19^ The BARIA cohort consists of individuals with obesity undergoing bariatric surgery with a BMI of ≥40 kg/m^2^ or ≥35 kg/m^2^ with obesity-related comorbidities including hypertension, T2D, obstructive sleep apnea, osteoarthritis, or dyslipidemia (primary forms excluded). Samples were further analyzed through tissue transcriptomics, untargeted plasma metabolomics, and fecal microbial DNA sequencing.

We analyzed 125 BARIA cohort patients who had both a blood sample drawn on the morning of their bariatric surgery and undergone biopsy of all aforementioned tissues. Among these 125, we divided patients by the presence of dyslipidemia or not, where dyslipidemia was defined by patients being on lipid lowering medications, or having ≥1 of the NHANES ATP III criteria (criteria are described in the **supplementary**).^20^

The research was conducted in accordance with the Declaration of Helsinki and had approval from the Ethics Committee of the Amsterdam University Medical Center. All study participants gave written informed consent prior to participation.

### Data availability

Data is available upon reasonable request through contacting the corresponding author.

### Sample collection

Baseline characteristics were obtained after an overnight fast on participant’s pre-operative visit. Participants collected stool samples on the day prior to their planned bariatric surgery. Upon arrival at the admission department for surgery, fecal samples were instantly frozen at -80°C. For plasma lipids, lipoproteins, and adipokines, we obtained venous blood samples on EDTA-coated tubes on the morning of bariatric surgery, with centrifugation at 4°C for 15 minutes, we isolated the plasma fraction and stored it at -80°C until further processing. Biopsies of SAT, VAT, liver, and jejunum were performed perioperatively and snap frozen in liquid nitrogen within 20 seconds of biopsy, ensuring minimal RNA breakdown. Biopsies were obtained within 2 hours of blood drawing.

### Plasma sample analysis

For measurement of total cholesterol, TG, HDL-C, Apo-I, ApoB, we used commercially available kits (DiaSys, WAKO) on an automated clinical chemistry analyzer (Selectra, Sopachem, Netherlands). We determined LDL-C with the Friedewald equation.^21^ We further investigated the concept of adipocyte dysfunction through measurement of adipokines and determined the plasma adiponectin-leptin ratio (A/L ratio) as a marker of dyslipidemia. Plasma adiponectin and leptin concentrations were measured using human Adiponectin/Acrp30 DuoSet^®^ ELISA (Y1065-05; R&D Systems, Minneapolis, USA) and human Leptin DuoSet^®^ ELISA (DY398-05; R&D Systems, Minneapolis, USA).

### RNA sequencing of tissue samples

Tissue transcriptomics methods are explained in detail in **Supplementary Methods**. Sample sizes for tissue transcriptomics were SAT n=116, visceral fat n=124, liver n=123, and jejunum n=118. Analyses were adjusted for potential confounders: age, sex, and sequencing batch. Next, we used the pathfindR R package version 2.3.0^22^ for pathway enrichment analysis, and KEGG database for biological pathway identification.

### Plasma metabolomics

Untargeted plasma metabolomics were done at Metabolon USA. We initially applied a binary approach (dyslipidemia present or not) and sought correlations with certain metabolites on the presence of dyslipidemia. Next, we applied continuous data correlation analysis by comparing continuous values of ApoAI, ApoB, triglycerides, cholesterol, HDL-C, and LDL-C from the day-of-surgery blood samples in correlation with found metabolites. Full details of metabolomics methodology in **Supplementary Methods**.

### Gut bacterial metagenomics

For gut bacterial metagenomics we only applied a binary approach (dyslipidemia being present or not) in relation to GM composition. DNA microbial metagenome sequencing data of 110 fecal samples were analyzed. Methods are detailed in the **Supplementary Methods**.

### Statistical analysis

Statistical analyses were performed using R v4.3.03^23^ and RStudio 2024.04.2^24^. Normally distributed data were described with mean (± standard deviation), non-normal with median (1^st^– 3^rd^ quartile). For categorical variable comparison we used Chi-squared testing for independence. We compared normally distributed continuous variables with student’s t test, and non-normal with Mann-Whitney U testing. Correlation of normally distributed continuous was done with Pearson’s correlation, and non-normal with Spearman’s correlation. Since we correlated multiple adipokines and plasma lipid, and lipoprotein levels, to reduce the false discovery rate (FDR), we applied the Benjamini-Hochberg correction to results of transcriptomic analysis and plasma biomarkers, repeated correlations were filtered for significant associations using the same adjustment method. We applied multiple linear regression modeling correcting for potential confounders in determining adipokine levels in dyslipidemia versus non-dyslipidemia. For transcriptomics analyses, we opted to report and interpret only the significant gene alterations with a ≥0.2 log_2_ fold change (DESeq2) for dyslipidemia status or ≥0.2 β-coefficient (MaAsLin) for continuous lipid associations. For all statistical comparisons and tests, we considered a statistical significance level (*p*) of <0.05 as significant.

## Results

A visual summary of the principal transcriptomic and metabolomic alterations associated with dyslipidemia and plasma lipid profiles is given in **Figure 1**.

**Figure 1.**
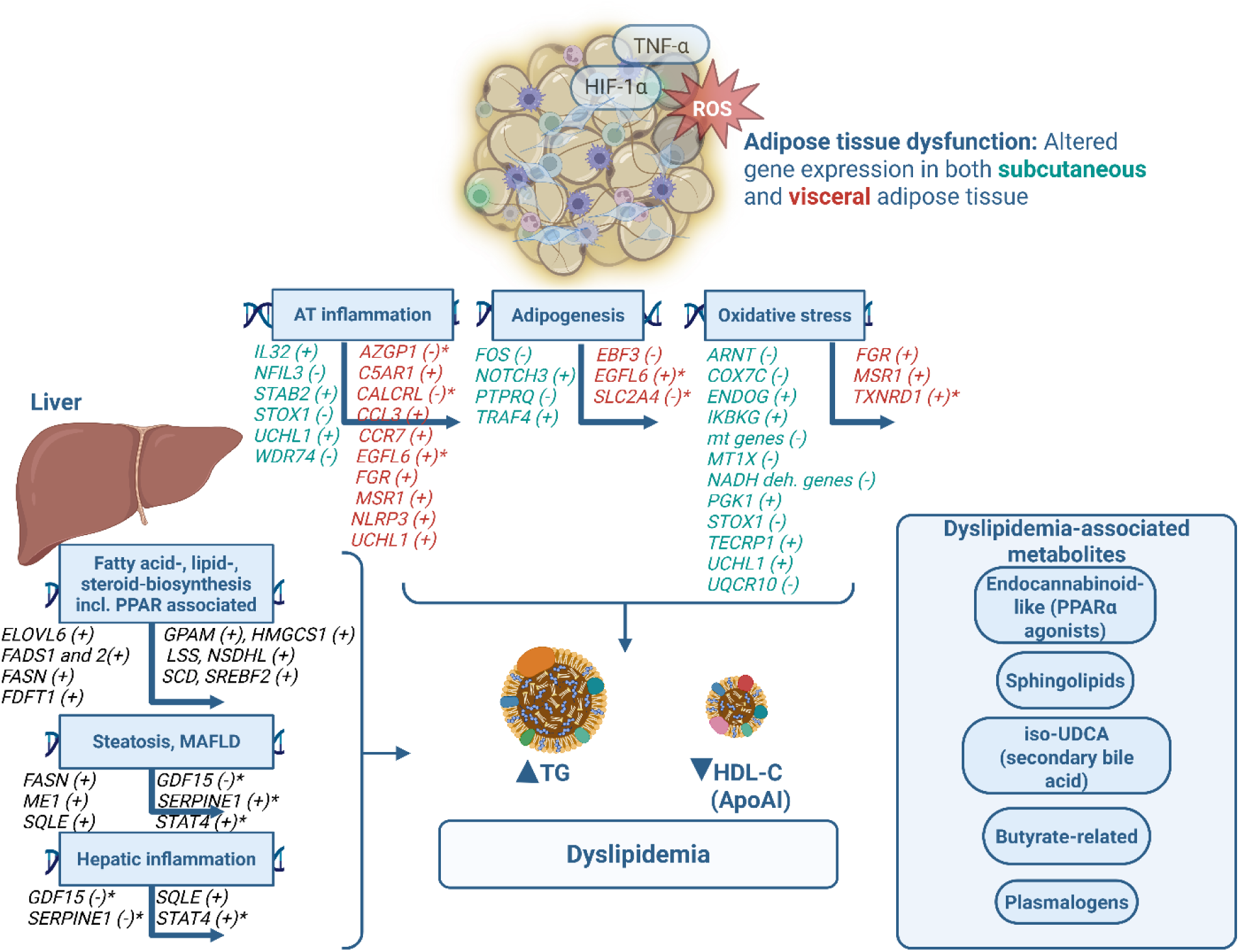
Summary of transcriptomic and metabolomic results. Genes expressed in SAT, VAT or liver with positive (+) or negative (-) associations with dyslipidemia or a deleterious lipid profile (elevated TG or decreased HDL-C or ApoAI). Genes described in detail in table S1 and S2. *These genes have an inverse relationship with HDL-C, meaning if positive (+), they were positively related to dyslipidemia (negative correlation with HDL-C), and if negative (-), they were negatively related to dyslipidemia (positive correlation with HDL-C). Created in BioRender. Zwartjes, M. (2025) https://BioRender.com/j4alzlj.

### Patients and plasma biomarkers

We included 125 patients, with a mean BMI of 39.5 kg/m^2^, and 80.8% being female. Patient characteristics are shown in **Table 1**. Dyslipidemia was present in 43 patients (34.4%), whereas 82 (65.6%) had a normal lipid profile.

**Table 1.** Patient Characteristics

|  | Total population<br>(n=125, 100%) | Dyslipidemia<br>(n=43, 34.4%) | No dyslipidemia<br>(n=82, 65.6%) | *p* |
| --- | --- | --- | --- | --- |
| **Demographic** |  |  |  |  |
| Female | 101 (80.8%) | 28 (65.1%) | 73 (89.0%) | <.001 |
| Age (years) | 44.3 ± 10.4 | 42.6 ± 10.6 | 42.3 ± 10.3 | .176 |
| **Anthropometric** |  |  |  |  |
| Weight (kg) | 115.0 ± 13.8 | 122.2 ± 14.4 | 112.4 ± 13.8 | .004 |
| BMI (kg/m<sup>2</sup>) | 39.5 ± 3.6 | 39.6 ± 3.1 | 39.4 ± 3.8 | .800 |
| Waist circumference (cm) | 121.4 ± 10.1 | 123.7 ± 10.8 | 120.1 ± 9.5 | .058 |
| Hip circumference (cm) | 130.7 ± 9.4 | 130.6 ± 9.4 | 130.8 ± 9.5 | .905 |
| Waist-to-hip ratio | 0.9 ± 0.1 | 0.9 ± 0.1 | 0.9 ± 0.1 | .056 |
| Body fat (kg) | 53.7 ± 8.4 | 54.6 ± 7.5 | 53.29 ± 8.9 | .470 |
| Systolic blood pressure (mmHg) | 122.0 ± 15.0 | 122.0 ± 15.0 | 121.0 ± 15.1 | .853 |
| Diastolic blood pressure (mmHg) | 79.1 ± 9.4 | 77.1 ± 8.6 | 80.1 ± 9.6 | .093 |
| Heart rate (bpm) | 73.9 ± 11.0 | 70.6 ± 9.9 | 75.7 ± 11.2 | .011 |
| **Clinical lab values at baseline** |  |  |  |  |
| ALP (U/L) | 83.8 ± 21.7 | 84.4 ± 21.0 | 83.5 ± 22.2 | .826 |
| GGT (U/L) | 28.0 (20.0–41.0) | 27.0 (19.5–41.0) | 28.0 (20.0–39.75) | .920 |
| AST (U/L) | 23.0 (20.0–28.0) | 25.0 (21.0–29.0) | 23.0 (20.0–27.5) | .144 |
| ALT (U/L) | 30.0 (22.0–38.5) | 29.0 (22.0–37.0) | 31.0 (22.0–39.3) | .609 |
| Ferritin (μg/L) | 85 (47.5–171.5) | 131 (81.0–196.0) | 75.5 (33.8–134.8) | .002 |
| CRP (mg/L) | 6.8 (3.5–9.0) | 5.0 (3.0–9.0) | 7.4 (4.0–9.0) | .160 |
| HbA<sub>1c</sub> baseline (%) | 5.41 ± 0.4 | 5.41 ± 0.3 | 5.42 ± 0.4 | .951 |
| **Baseline mixed meal test** |  |  |  |  |
| Fasting glucose (mmol/L) | 5.67 ± 0.6 | 5.72 ± 0.5 | 5.64 ± 0.7 | .458 |
| Fasting insulin (mIU/L) | 82.65 ± 47.1 | 84.87 ± 36.3 | 81.52 ± 52.0 | .709 |
| C-peptide (ng/mL) | 0.76 ± 0.3 | 0.82 ± 0.3 | 0.73 ± 0.3 | .124 |
| HOMA-1 IR | 2.79 (1.80–3.98) | 3.09 (2.29–4.27) | 2.37 (1.63–3.86) | .123 |
| HOMA-2 IR | 1.49 (0.98–2.09) | 1.69 (1.35–2.16) | 1.29 (0.97–2.02) | .068 |
| **Liver histology** |  |  |  |  |
| No steatosis | 58 (47.2%) | 17 (39.5%) | 41 (51.2%) |  |
| MASLD | 49 (39.8%) | 19 (44.2%) | 30 (37.5%) |  |
| MASH | 16 (13.0%) | 7 (16.3%) | 9 (11.2%) |  |
|  |  |  |  | .434<sup>†</sup> |

Data expressed as mean ± standard deviation (SD) or as median (1<sup>st</sup>–3<sup>rd</sup> quartile); <sup>†</sup>p-value of $\chi^2$ -test for independence; ALP: alkaline phosphatase; ALT: alanine transaminase; AST: aspartate transaminase; CRP: C-reactive protein; GGT: $\gamma$ -glutamyltransferase; HOMA IR: homeostatic model assessment for insulin resistance. For calculating HOMA, we used “HOMA-1= (fasting plasma glucose [mmol/L] x fasting plasma insulin [ $\mu$ U/mL] x) / 22.5”,<sup>69</sup> and to acquire HOMA-2 values,<sup>70</sup> we used the online available HOMA-2 calculator software;<sup>71</sup> MASLD: metabolic dysfunction-associated steatotic liver disease; MASH: metabolic dysfunction-associated steatohepatitis.

Complete lipid and lipoprotein concentrations are seen in **Table 2**. Patients with dyslipidemia had lower HDL-C, and ApoAI, and higher TGs, ApoB, whereas LDL-C did not differ in patients with and without dyslipidemia. With adipokines, individuals with dyslipidemia did not have a difference in adiponectin concentration (adjusted *p* = .072), and, unexpectedly, leptin concentration was lower in individuals with dyslipidemia (adjusted *p* = .010). A/L ratio between the groups did not differ (adjusted *p* = .160).

**Table 2.**
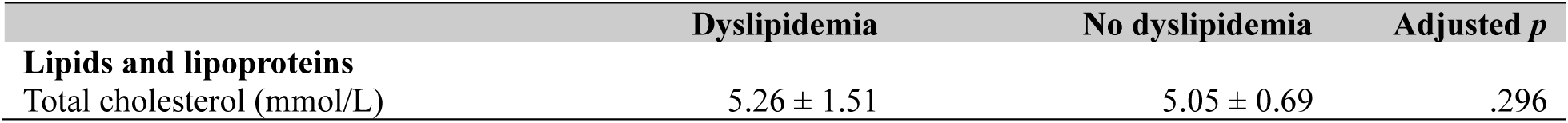
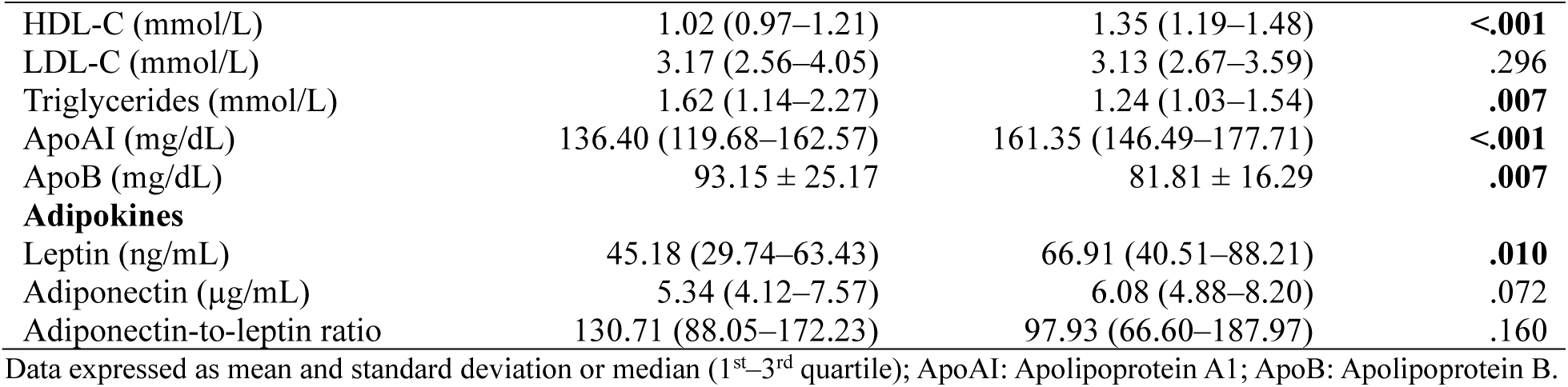
Plasma Biomarkers on Day of Surgery

As mentioned, patients with dyslipidemia had lower leptin concentrations. Since patient characteristics of the groups showed differences in gender, age, and body weight (**Table 1**), we performed additional multiple linear regression analyses to account for these potential confounders: After adjustment, leptin did not differ between dyslipidemia and non-dyslipidemia patients (multiple linear regression *p*-value after adjustment: *p* = 0.057 for leptin). We did not include ferritin as a confounder in the model due to a lack of evidence in literature substantiating the direct influence of ferritin on leptin concentration.

Correlation analysis of adipokines with lipids or apolipoproteins is seen in **Figure 2** and **3**: Leptin showed weak positive correlation with HDL-C (Spearman’s ρ = 0.24, *p* = <0.01; **Figure 2A**) and its primary apolipoprotein ApoAI (Spearman’s ρ = 0.28, *p* = <0.01; **Figure 2B**). Adiponectin showed moderate positive correlation with plasma HDL-C (Spearman’s ρ = 0.30, *p* = <0.01; **Figure 3A**), weak positive correlation with ApoAI (Spearman’s ρ = 0.20, *p* = 0.03; **Figure 3B**) and weak negative correlations with triglycerides (Spearman’s ρ = –0.19, *p* = 0.03; **Figure 3C**), and ApoB (Spearman’s ρ = –0.20, *p* = 0.03; **Figure 3D**). Correlation of adipokines and other lipids or apolipoproteins was not significant. However, correlation analysis of adipokines with insulin resistance (HOMA-1 IR) revealed a moderate inverse relationship between adiponectin and HOMA-1 IR (Spearman’s ρ = –0.45, *p* = <0.001, **Figure S2**), but no correlation was found between HOMA-1 IR and leptin (Spearman’s ρ = 0.08, *p* = 0.63).

**Figure 2.**
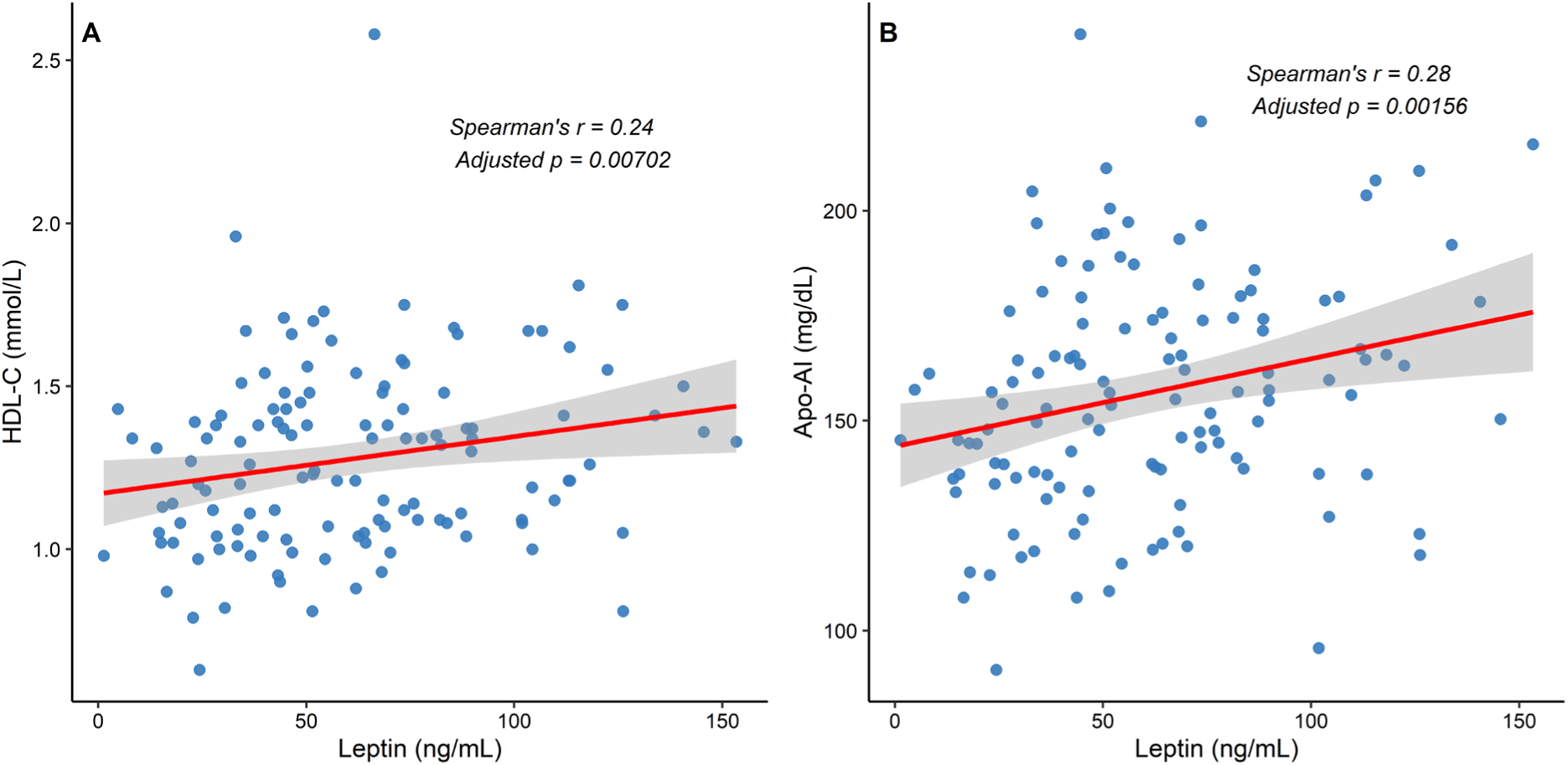
Correlation plots of leptin with HDL-C and ApoAI with lines of best fit and confidence interval. Spearman’s rank correlation (r), and its Benjamini-Hochberg adjusted p-value (*p*), only significant correlations depicted.

**Figure 3.**
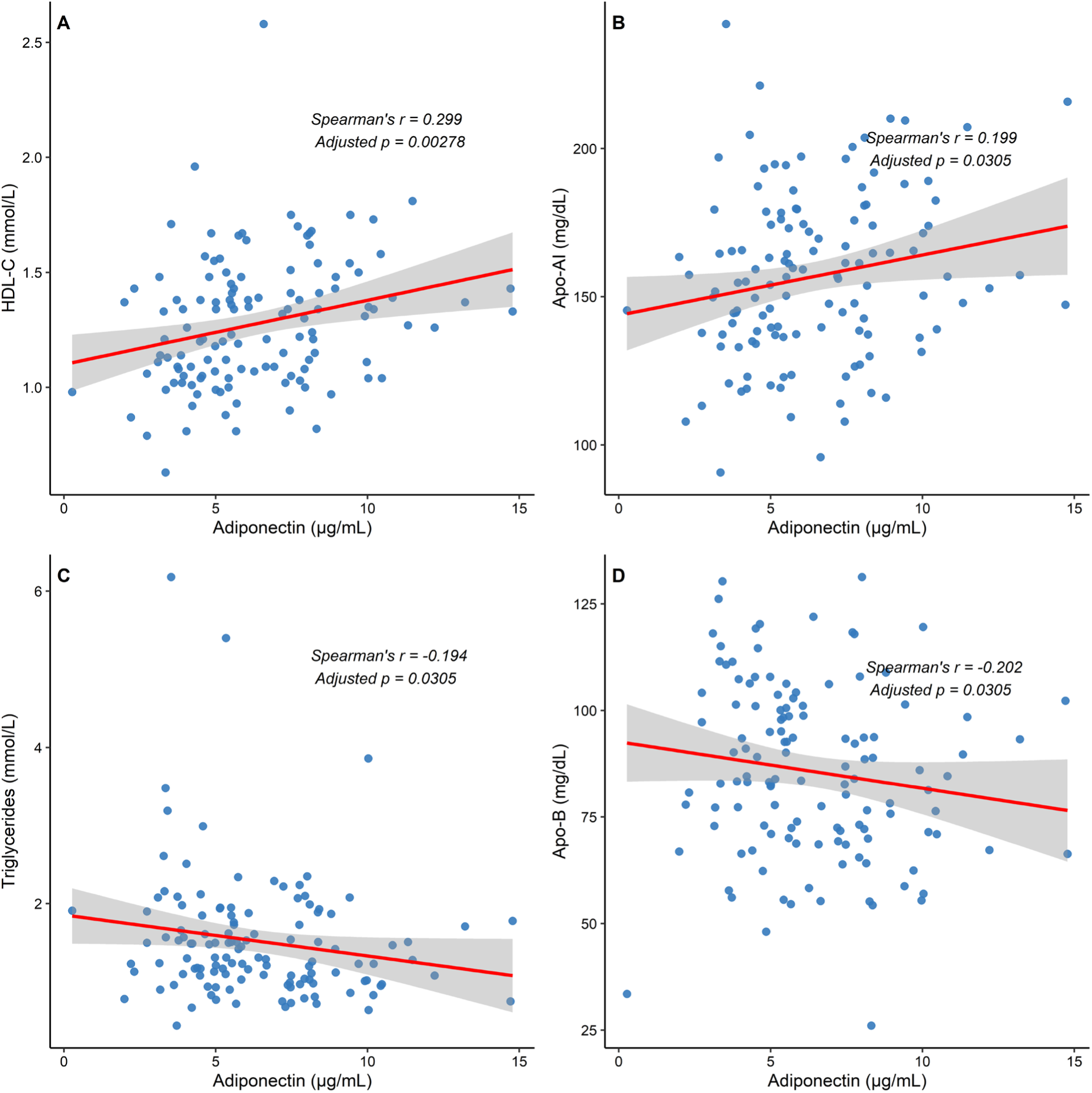
Correlation plots of adiponectin with lipids and apolipoproteins with lines of best fit and confidence intervals. Spearman’s rank correlation (r), and its Benjamini-Hochberg adjusted p-value (*p*), only significant correlations depicted.

### Transcriptomics

To determine differential gene expressions in patients with and without dyslipidemia, we correlated transcriptomics of SAT, VAT, jejunum, and liver tissues with dyslipidemia status or continuous lipid and lipoprotein concentrations. This revealed a large number of genes of which gene expressions correlated to plasma TG, HDL-C and ApoAI, but no correlations were found with total cholesterol, LDL-C or ApoB. **Figure 1** shows an overview, and a description of identified genes with functional implications, can be found in **Table S1** and **S2**.

In SAT (116 patients, 40 dyslipidemia), genes were up- or downregulated in association with dyslipidemia status and plasma TG. As outlined in **Figure 1** (and **Table S1** and **S2**), we found associations of genes known to be important in not only lipid metabolism, but also in (adipose tissue) inflammation, adipogenesis, and oxidative stress. In SAT, when comparing dyslipidemia status with gene expression, *IL32* was upregulated in dyslipidemia, and *FOS*, *NFIL3*, *WDR74* downregulated (**Figure S3A**). In males with dyslipidemia, *STAB2* and *TRAF4* were upregulated, and *MT1X* downregulated. Genes related to inflammation that correlated with plasma TG or dyslipidemia status were: *IL32*, *NFIL3*, *STAB2*, *STOX1*, *UCHL1*, and *WDR74*. Genes from oxidative stress (ROS) pathways were upregulated (*ENDOG*, *TECRP1*, *UCHL1*) or downregulated (*MT1X*, *mitochondrial [mt] genes [predominantly downregulated]*, *STOX1*). Genes important for adipogenesis were also identified (*FOS*, *NOTCH3*, *PTPRQ*, *TRAF4*) and correlated with dyslipidemia status or plasma TG. **Figure 4** illustrates some of enriched pathways in SAT identified by KEGG analysis and PathfindR in relation to plasma TG concentration: These include Notch signaling pathway (4.6-fold enrichment, including *NOTCH3* upregulation), Hypoxia-inducible factor 1 (HIF-1) signaling (2.8-fold enrichment, upregulated *PGK1*, downregulated *ARNT*, *CREBBP*, *EIF4EBP1*, *EP300*, *RPS6*). We also found enrichment of oxidative phosphorylation pathways (*OXHPOS* genes, 2.6-fold enrichment) illustrated in **Figure 5**; including downregulation of complex I (NADH dehydrogenase genes: *NDUFA4*, *NDUFA7*, *NDUFA12*, *NDUFB4*), complex III (cytochrome bc1 complex; *UQCR10*), and complex IV (*COX7C*) genes. We found alteration of chemical carcinogenesis and ROS production pathways (1.9-fold enrichment, **Figure S4**) with upregulated (*IKBKG*) and downregulated genes (4 NADH dehydrogenase genes, as well as *UQCR10*, *COX7C* and *ARNT*).

**Figure 4.**
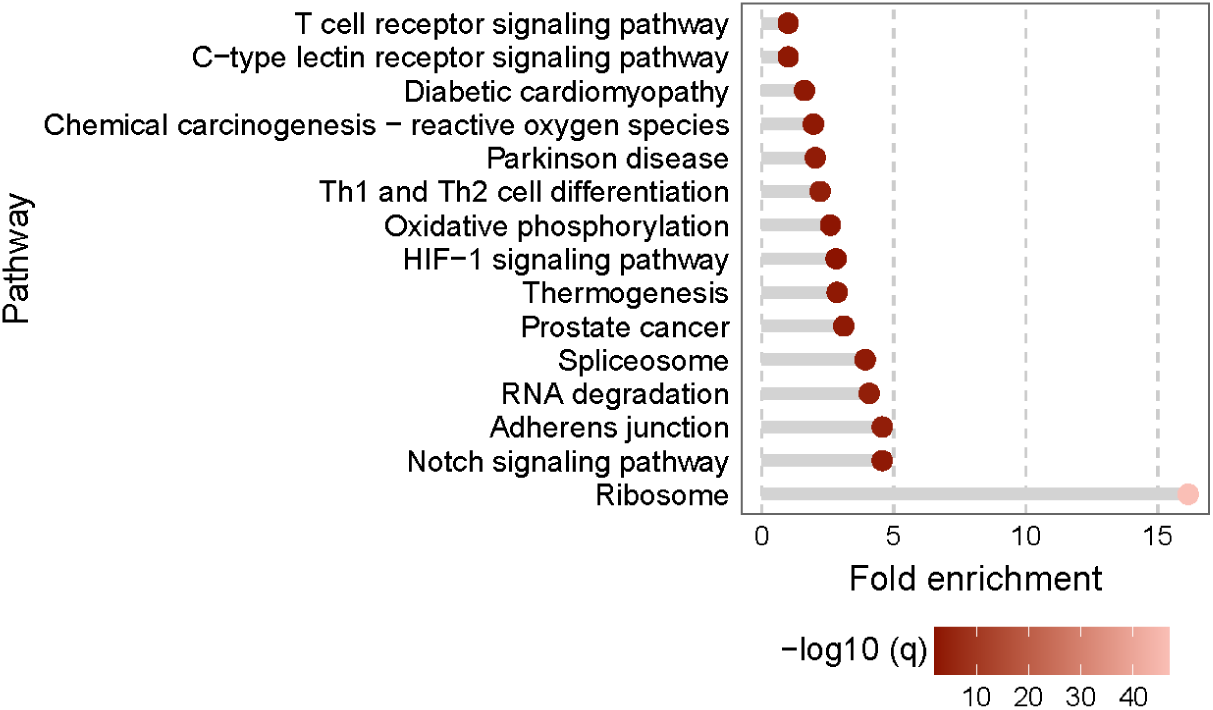
Dot plot of differential gene expression of SAT in relation to plasma TG through KEGG pathway analysis. X-axis represents pathway fold enrichment. Dot color is the –log10(q) value, where the q-value (adjusted *p*-value) indicates significance of pathway enrichment. Using –log10(q) transforms small q-values into larger positive numbers.

**Figure 5.**
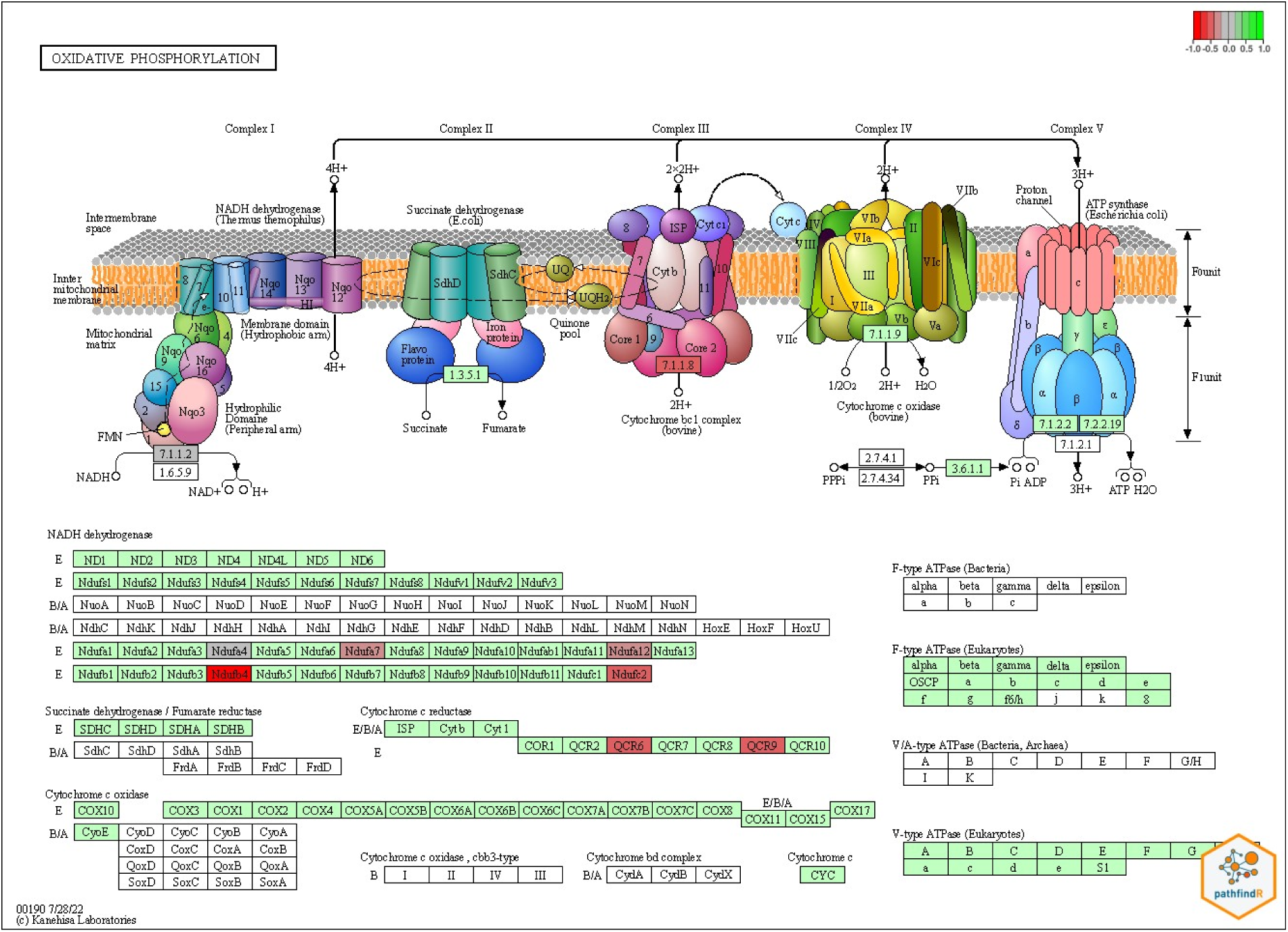
Oxidative phosphorylation (OXPHOS genes): Relative gene expression (pathfindR) of SAT genes of the oxidative phosphorylation pathway in relation to plasma triglycerides. Color coding from –1 (red) downregulated, to +1 (green) upregulated genes, –1 and +1 signifies a log2 fold change.

In VAT (124 patients, 43 dyslipidemia), we found gene correlations with plasma TG, HDL-C, and ApoAI. In dyslipidemia we found downregulated *EBF3* and upregulated *APOA4* (**Figure S3B**, **Table S1**). In VAT, altered expressions of genes with functions in inflammation were: *AZGP1*, *C5AR1*, *CALCRL*, *CCL3*, *CCR7*, *EGFL6*, *FGR*, *MSR1*, *NLRP3*, and *UCHL1*. In oxidative stress: *FGR*, *MSR1*, and with minor positive correlation: *TXNRD1*. In adipogenesis or adipocyte differentiation: *EBF3*, *EGFL6*, *SLC2A4*. In relation to plasma TG, pathfindR KEGG analysis revealed enrichment of several pathways including ribosome (9.2-fold), NOD-like receptor signaling (8.0-fold), lipid and atherosclerosis (9.8-fold, including *BCL2L1*, *CCL3* and *NLRP3* upregulation).

Liver transcriptomics (123 patients, 41 dyslipidemia) and pathfindR KEGG analysis found not only genes involved in *de novo* lipogenesis or lipid metabolism (steroid biosynthesis; 60.0-fold enrichment, unsaturated fatty acid biosynthesis; 46.7-fold, PPAR signaling; 15.2-fold), but also genes associated with liver steatosis or inflammation, these genes were *FASN*, *FDFT1*, *GDF15*, *ME1*, *SERPINE1*, *SQLE*, *STAT4*. Genes are described in **Figure 1**, and in detail in **Table S1** and **S2**.

In jejunum (118 patients, 41 dyslipidemia), only two genes were found to be differentially expressed in patient with versus without dyslipidemia: *HOXC6* and *LCN2* (**table S1**), both upregulated in dyslipidemia. No correlation was found with continuous lipid or apolipoprotein concentration.

## Metabolomics

Several plasma metabolites associated with dyslipidemia and are shown in **Figure 6**. Palmitoyl ethanolamide (PEA) and oleoyl ethanolamide (OEA), both endocannabinoid-like lipid metabolites and PPARα agonists, positively associated with dyslipidemia. The GM-derived secondary bile acid, isoursodeoxycholate (isoUDCA) positively correlated to dyslipidemia. Sphingolipids and plasmalogens had negative correlations. We discerned a total of 234 metabolites correlating to plasma lipids, lipoprotein, or adipokines (**Figure S5** and **S6**). We found mainly positive correlations with plasma lipids, including monoacylglycerols and lipid derivatives. Some lipid metabolites, however, showed negative correlations, including butyrate-related metabolites, sphingolipids and plasmalogens which negatively correlated to TG. Several metabolites correlated with adipokines, including butyrates which had negative correlations with leptin.

**Figure 6.**
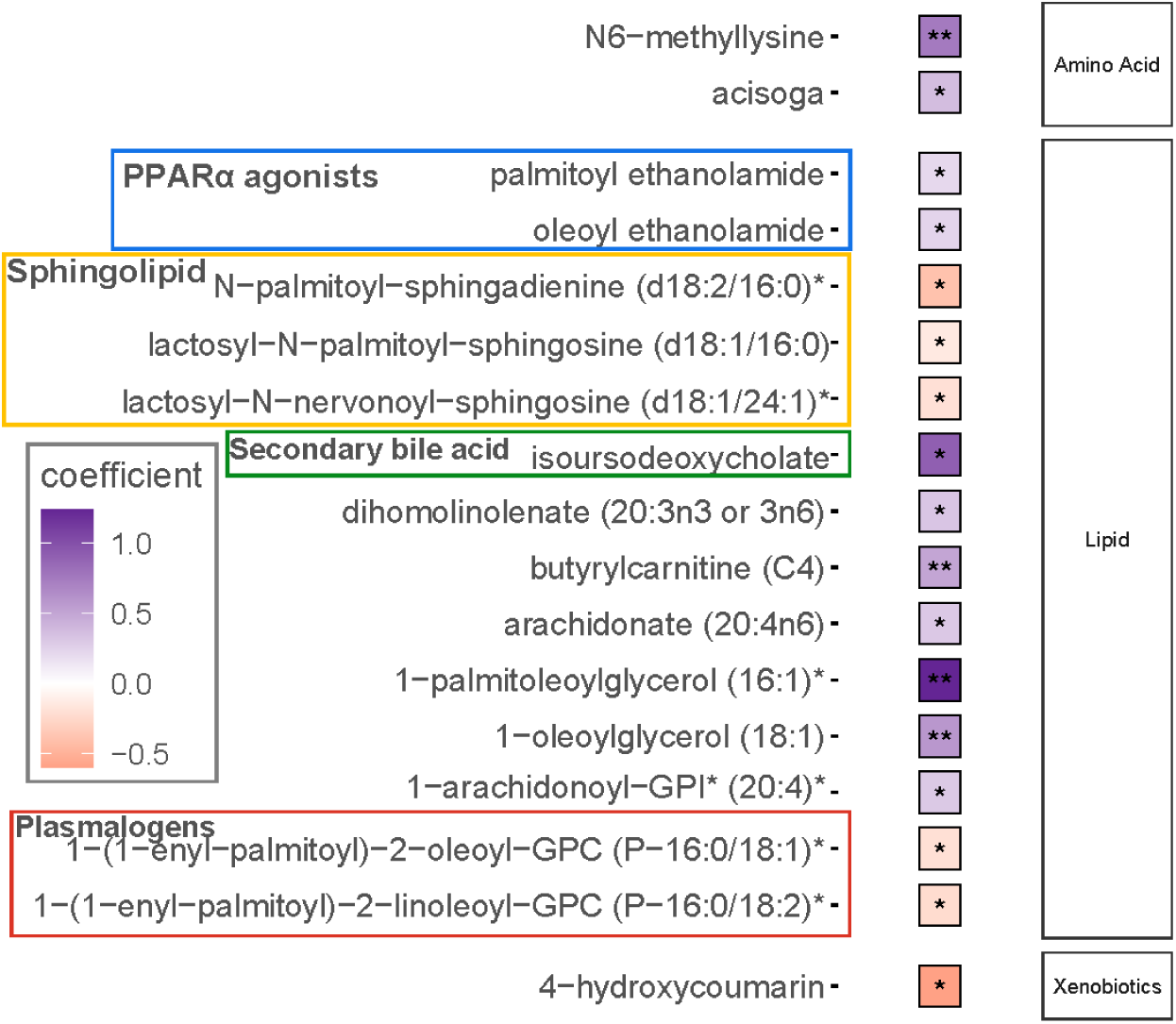
Metabolites in dyslipidemia. The coefficient refers to the MaAsLin2 correlation coefficient to dyslipidemia status; *: *p*<.05; **: *p*<0.01; ***: *p*<.001.

## Gut microbiota analyses

Only a modest difference in gut microbiota composition (relative abundance) between patients with and without dyslipidemia was found, results of which are detailed in **Supplementary Results** and **Figure S7–S9**). On a bacterial order level, in patients with dyslipidemia, we found more species of the order *Lachnospirales* and of the order *Peptostreptococcales*, both part of the Firmicutes phylum. In patients without dyslipidemia, *Oscillospirales* (also within the Firmicutes phylum), *Bacteroidales* and *Lactobacillales* predominated (**Figure S9**).

## Discussion

We investigated 125 individuals with obesity of which about one third had dyslipidemia. Individuals with obesity and dyslipidemia had higher TG and ApoB and lower HDL-C and ApoAI, a typical pattern of dyslipidemia seen in obesity.^25^ Unexpectedly, we found no difference in adiponectin and even lower leptin in patients with dyslipidemia, A/L ratio also did not differ. This is in contrast to earlier studies highlighting leptin as pro-atherogenic and detrimental for health and adiponectin as protective (insulin sensitizing, anti-atherogenic).^9,14,16^

Next, in contrast to our hypothesis of the predominant role of VAT in dyslipidemia, our transcriptomics revealed that both VAT *and* SAT had altered transcriptome in relation to dyslipidemia or plasma lipids, and notably, SAT had most numerous gene alterations. We found that genes were up- or downregulated in relation to plasma TG, HDL-C, and ApoAI, rather than cholesterol or ApoB. Since previously mainly VAT associated with cardiometabolic disease,^10–12^ our finding of altered SAT transcriptome in dyslipidemia underscores an underestimated role of SAT in dyslipidemia pathogenesis. We identified 3 recurring pathways in AT in relation to dyslipidemia or lipid profiles: (1) AT inflammation, (2) adipogenesis, and (3) oxidative stress.

AT inflammation has been described as a factor in cardiometabolic disease development.^6,13^ We identified several altered inflammatory pathways in relation to dyslipidemia: In SAT, pro-inflammatory cytokine gene *IL32* was upregulated, and the TGF-β co-activator gene, *WDR74*, downregulated; TGF-β is essential for adipogenesis and immune homeostasis, and impaired TGF- β signaling contributes to inflammatory pathologies.^26^ In SAT of patients with dyslipidemia, *NFIL3* was downregulated, *NFIL3* produces E4bp4, a transcription factor maintaining anti-inflammatory AT homeostasis through IL-10 mediated M2 macrophage polarization and promotion of regulatory T cells.^27,28^ *NFIL3* downregulation in dyslipidemia possibly reflects a shift towards a more pro-inflammatory environment. We found strong *STAB2* upregulation in males with dyslipidemia; *STAB2* is a scavenger receptor indirectly regulating inflammation through affecting clearance of pro-inflammatory ligands and a murine study showed that *STAB2* knockout reduced atherosclerosis.^29^ In both SAT and VAT, *UCHL1* expression correlated to elevated TG, in line with previous reports that *UCHL1* is upregulated in obese AT and increases inflammation through degradation of RXRα and inhibition of LXR/RXR’s anti-inflammatory effects.^30^ In VAT, pro-inflammatory *CCL3* positively correlated with plasma TG, and *CALCRL* with HDL-C.

*CALCRL* encodes adrenomedullin receptor with anti-inflammatory effects through NF-κB inhibition, and reduces inflammatory cytokines and CRP, whereas CCL3 is released by inflamed AT in obesity, working as a chemoattractant for monocytes and macrophages partly through NF- κB.^31,32^ NF-κB signaling pathway is likely an important driver of AT inflammation and ROS production during overnutrition and metabolic dysregulation.^33^ In VAT, *AZGP1*, correlated positively with HDL-C and adiponectin, AZGP1 is an adipokine increasing lipolysis, AT browning, and insulin sensitivity by, amongst others, suppression of pro-inflammatory pathways involving NF-κB, IL1β and IL-6.^34^ In VAT, *CCR7* had positive association with plasma TG, reflecting its possible role in coordinating immune cell trafficking, sustaining AT inflammation.^35^ Complement pathways were also implied in our findings; we found *C5AR1* upregulation in VAT in relation to TG, this gene encodes the receptor for complement component C5a, which promotes inflammation through recruitment of M1 macrophages.^36^ Lastly, we found positive correlation between *NLRP3* and TG, its transcription leads to inflammasome activation and inflammatory cytokine IL-1β and IL-18 production in AT.^37^

Adipogenesis emerged as the next key transcriptomic pathway. In SAT, *FOS*, a critical regulator of pre-adipocyte differentiation through its interaction with AP-1,^38^ was downregulated in dyslipidemia, and *NOTCH3*, which promotes early adipogenesis and PPARγ expression,^39^ positively correlated with plasma TG. In SAT, *PTPRQ*, another regulator of adipogenesis via the PI3K-PKB/Akt pathway,^40^ correlated negatively with TG. In males with dyslipidemia, *TRAF4* was upregulated in SAT, consistent with its function as adipogenesis checkpoint regulator.^41^ In VAT, *EGFL6* expression negatively correlated with HDL-C; a study analyzing AT in children with obesity found *EGFL6* as a key secreted extracellular matrix protein with upregulation during adipocyte differentiation and remodeling, and strong associations with adipose hypertrophy, macrophage infiltration, hs-CRP, and insulin resistance.^42^ Its negative correlation with HDL-C, a protective lipoprotein, might reflect *EGFL6’s* contribution to dyslipidemia through altered adipocyte development. In VAT, *EBF3*, a transcription factor for adipogenesis and brown adipocyte development, was downregulated in dyslipidemia, while *SLC2A4*, another gene of adipogenesis, had positive correlation with HDL-C. Adipogenesis regulation is complex, and our findings indicate both up- and downregulated adipogenesis genes in relation to dyslipidemia.

Thirdly, we repeat earlier findings^43^ of upregulated oxidative stress genes in dysfunctional AT: we identified genes of oxidative phosphorylation, ROS production and HIF-1. *UCHL1* expression in SAT and VAT had positive relation with TG, *UCHL1* modulates oxidative stress response through LXR/RXR signaling.^30^ Autophagy is essential for response to oxidative stress,^44,45^ in SAT we found strong positive relation between *TECRP1* and TG, this is relevant because *TECRP1* produces one of the main autophagy tethering proteins facilitating autophagosome-lysosome fusion to regulate ROS. *ENDOG*, the gene for a mitochondrial nuclease involved in autophagy regulation and oxidative stress response,^46^ also positively correlated with plasma TG in SAT. Another gene counteracting oxidative stress, *STOX1*, correlated negatively with plasma TG in SAT. In males with dyslipidemia, *MT1X* in SAT was downregulated, this is a metallothionein protecting against oxidative stress through free radical scavenging,^47^ downregulation might indicate reduced oxidative stress resilience in AT dysfunction. In VAT, *FGR* associated positively with plasma TG, *FGR* is a downstream effector of ROS signaling and is essential for proinflammatory M1-like macrophage polarization in AT, driven by mitochondrial ROS and complex II activation.^48^ Next, our PathfindR KEGG analysis found alteration of several oxidative phosphorylation complex genes in relation to plasma TG, but also genes of HIF-1 pathways; Hypoxia inducible factor, as the name implies, responds to hypoxia and is either HIF-1α or HIF-1β. Under normoxia HIF-1α is readily degraded, under hypoxia, HIF-1α persists, heterodimerizes with HIF-1β, and binds hypoxia responsive element (HRE) in DNA, causing gene expression.^49^ We found upregulated *PGK1* in SAT associated with plasma TG; *PGK1* is induced under hypoxic conditions by HIF-1α signaling and can inhibit angiogenesis,^50^ supporting a concept of dysfunctional AT where its expansion exceeds vascularization potential. Another indicator of reduced angiogenesis in dysfunctional AT is our observed *ARNT* downregulation in patients with elevated TG; *ARNT* (termed HIF-1β) normally *promotes* angiogenesis through transcription complex formation with HIF-1α.^51^. Downregulation of oxidative phosphorylation genes in relation to TG (particularly of mitochondrial respiratory chain complexes I, III, and IV), indicates altered mitochondrial energy metabolism and our found downregulation of these genes could highlight a dysfunctional AT with reduced ability of inducing mitochondrial energy metabolism at the time of biopsy (fasted state). In VAT we found altered expressions oxidative stress genes, *FGR* and *MSR1* (both positive correlation with TGs), and *TXNRD1* (minor negative correlation with HDL-C). *TXNRD1* activity has been shown to protect from oxidative stress and ROS release.^52^

Hepatic transcriptome confirmed alterations of genes involved in lipid metabolism and *de novo* lipogenesis in relation to plasma TG but also highlighted genes with roles in hepatic steatosis or inflammation (MAFLD or MASH) including: *FASN*, *GDF15*, *ME1*, *SERPINE1*, *SQLE*, *STAT4*. *SQLE*, the gene for squalene epoxidase is the second rate-limiting enzyme of cholesterol synthesis and has been found to contribute to hepatic steatosis and MASH.^53^ *SLC2A8*, encoding a fructose-glucose transporter GLUT8 also described in liver steatosis^54^. The lack of correlation between LDL-C related genes (including *HMGCR*, *LDLR*, *LDLRAP1*), might be explained by the fact that in obesity, dyslipidemia is less attributable to LDL-C, and rather by increased TG and reduced HDL-C.

Our metabolomics in dyslipidemia found correlation with palmitoyl ethanolamide (PEA) and oleoyl ethanolamide (OEA), both endocannabinoid-like lipid metabolites and PPARα agonists implicated not only in energy- and lipid metabolism, but also in inflammation and cardiovascular disease.^55,56^ OEA is likely involved in lipid metabolism through lipolysis enhancement,^57^ but is also described providing crosstalk between intestinal microbiota and intestinal homeostasis, where its malfunction is associated with enteropathies, obesity, and chronic inflammation.^58^ Studies have shown that PEA has anti-inflammatory action and can attenuate atherosclerotic plaque formation, decrease fat accumulation and lower plasma lipids.^59–61^ Next, we found positive correlation between isoUDCA and dyslipidemia; isoUDCA is a GM-derived bile acid, and a recent study already postulated its influence on lipid metabolism.^62^ Sphingolipid metabolism also appears to influence lipid metabolism and cardiometabolic disease,^63^ we found negative relation between sphingolipid metabolites and dyslipidemia. Butyrate-related metabolites, besides being endogenously produced from normal lipid metabolism, are short chain fatty acids (SCFAs) produced by the GM. SCFAs are increasingly investigated for their protective or anti-inflammatory roles and ability to reduce dyslipidemia, atherosclerosis and CVD risk,^64,65^ however, in our analysis these metabolites showed negative correlations with HDL-C and ApoAI; thus the beneficial aspect is less clear here but may indicate a regulatory function through HDL-C. Plasmalogens negatively associated with plasma TG, and a previous study has identified reduced plasmalogens in cardiometabolic disease.^66^ Plasmalogens appear to regulate thermogenic fat activation by induction of mitochondrial fission,^67^ potentially increasing fatty acid oxidation and reduce plasma TG.

Only modest differences in relative abundance were found between the gut metagenome of patients with and without dyslipidemia. Contrary to previous research linking SCFA-producing bacteria primarily to metabolic health,^64,65^ we found orders of SCFA producing bacteria in the gut metagenome of patients with (*Lachnospirales*) and without (*Oscillospirales*, including butyrate-producers) dyslipidemia, suggesting a more nuanced role of SCFA-producing bacteria in lipid metabolism. In patients without dyslipidemia, *Bacteroidales* and *Lactobacillales* orders predominated; *Bacteroidales* include the order *Bacteroidetes*, associated with lower MI risk,^68^ of which dyslipidemia is a major risk factor.

Our study carries limitations, including investigating dyslipidemia only in obesity and a predominance of female participants, affecting generalizability to the general population. Furthermore, the cross-sectional design and nature of transcriptomics limits identification of causal relationships.

In conclusion, our study provides a well-characterized cohort with multi-omics profiling for novel insights into dyslipidemia in obesity. In line with previous studies,^10–12^ our findings confirm the role of VAT in dyslipidemia but also highlight an underappreciated role of SAT in lipid metabolism and dyslipidemia. Increased AT inflammation, oxidative stress, and alterations of adipogenesis in both SAT and VAT appear to be the main pathways characterizing AT health versus dysfunction. Metabolomics revealed a complex interplay of metabolites in relation to lipids and dyslipidemia, highlighting potential roles of endocannabinoid-like metabolites, regulatory functions of GM-derived isoUDCA and butyrate-related metabolites. Future studies incorporating individuals from all BMI groups could further help identify the contribution of AT to dyslipidemia. And further studies will be required to clarify how our identified metabolites can be used prognostically, and how the identified AT dysfunction pathways can be altered for treating dyslipidemia and cardiovascular disease.

## Acknowledgements

The authors thank the BARIA study team for their dedicated assistance in participant recruitment and sample collection, as well as the entire bariatric surgery team from Baria Nederland in Spaarne Gasthuis Hoofddorp for allowing us to take obtain tissue samples during surgery. Additionally, we thank the Experimental Vascular Medicine lab department for their expert help with sample processing and measurements.

## Supplementary

### Supplementary Methods Study design and population

We defined dyslipidemia according to the National Cholesterol Education Program Adult Treatment Panel III (NHANES ATP III) criteria: total cholesterol ≥ 6.22 mmol/L (≥ 240 mg/dL), triglycerides > 2.26 mmol/L (≥ 200 mg/dL), LDL-C ≥ 4.14 mmol/L (≥ 160 mg/dL), and HDL-C < 1.04 mmol/L (< 40 mg/dL).^20^

#### Tissue transcriptomics

RNA was isolated from biopsied tissues by TriPure Isolation Reagent (Roche, Basel; Switzerland), Lysing Matrix D tubes and FastPrep-24^TM^ instrument (MP Biomedicals, Irvine, CA, USA) and through 20 seconds, 4.0 m/s homogenizations until no visible tissue remained, with samples kept on ice between homogenization cycles. Chloroform purification was done in with centrifugations at 4°C in phase separation tubes (Phase Lock Gel™, Quantabio LLC, Beverly, MA, USA). RNA purification was continued by an RNeasy MinElute cleanup kit (QIAGEN Benelux B.V., Venlo, Netherlands). For RNA quality assessment we used a bioanalyzer device (Agilent, Santa Clara, CA, USA) and purity measured using a NanoDrop^TM^ spectrophotometer (Thermo Fisher Scientific Inc, Waltham, MA, USA). RNA sequencing was performed at Novogene (Nanjing, China) using the Illumina HiSeq (Illumina Inc., San Diego, California). We performed quality trimming of reads with trimmomatic v0.38 (options: HEADCROP: 6, SLIDINGWINDOW: 4:15, and MINLEN: 50).^72^ Reads were mapped with Kallisto v0.46.0 against the GRCh38 human genome assembly (options --bias, -b 100, and --rf-stranded).^73^ We determined differential gene expression among patients with and without dyslipidemia with DESeq 2 v1.40.2 R package,^74^ and with continuous variables with MaAslin2 v1.14.1.^75^ As not all sequencing returned data of high-enough quality, some samples were excluded. SAT (n=116) data had some samples with smaller in library size than the rest (**Figure S1A**), and these samples with minor library size would disrupt our subsequent analysis through exertion of a dominant influence on the variation within the dataset as in a principal component analysis (PCA), axes 1 and 2 would account for >67% of the total variation observed between sample (**Figure S1B**), and these samples would therefore be responsible for around 67%, of the information contained in the dataset. Subsequently, removal of these samples for better interpretation was performed (**Figure S1C**). VAT (n=124), liver (n=123), and jejunum (n=118) transcriptomics had robust library size and could be analyzed without omitting samples.

**Figure S1.**
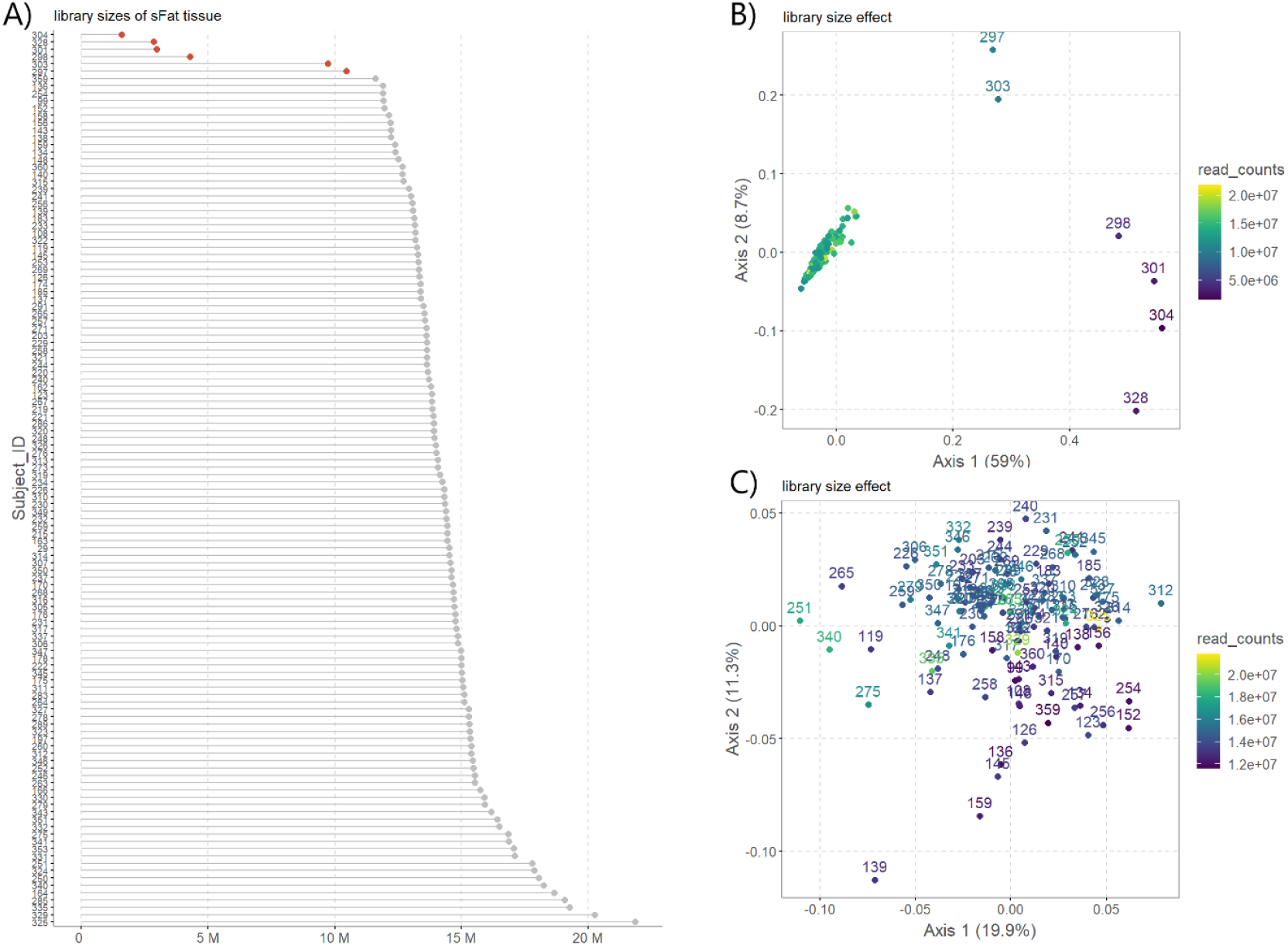
Library size of subcutaneous adipose tissue (SAT) for transcriptomics; A. SAT total library sizes; B. PCoA analysis of SAT samples; C. PCoA analysis of SAT samples after removal of samples with significantly smaller library sizes.

#### Plasma metabolomics

Plasma EDTA samples drawn from the day of surgery and were analyzed by METABOLON (Morrisville, NC, USA) using ultra high-performance liquid chromatography (UHPLC) and tandem mass spectrometry (LC-MS/MS) with an untargeted approach. Data was filtered initially to ensure less than 25% missing values for any metabolite in any of the samples. Subsequently, log transformation was applied to the raw signal values of all measured metabolites throughout the entire dataset. Of the 1346 metabolites identifiable by METABOLON, 998 metabolites were utilized in analysis after normalization and imputation. Non-annotated metabolites and those that derived from medications were excluded.

#### Gut bacterial metagenomics

Fecal samples provided, n = 110, were swiftly stored at -80°C. Aggregate bacterial metagenomic DNA was isolated out of 100 mg fecal matter with a modified IHMS protocol Q.^76^ Fecal material was placed into ASL buffer (Qiagen Benelux B.V., Venlo, Netherlands) inside Lysing Matrix E tubes (MP Biomedicals). Cells were lysed through a combination of mechanical disruption and heat treatment: vortexed for 2 minutes, then subjected to two cycles of heating at 90°C for 10 minutes each, and three rounds of mechanical homogenization with bead-beating at 5.5 m/s for 60 seconds using the MP Biomedicals™ FastPrep-24™ system. Supernatant with bacterial DNA was sampled after two centrifugations at 4°C, supernatants from both centrifugation steps were combined, and a 600 µL portion of each specimen underwent purification using a protocol of human DNA analysis with the QIAamp DNA Mini QIAcube Kit (QIAGEN Benelux B.V., Venlo, Netherlands). Elution was done by using a 200 mL AE buffer (0.5 mM EDTA with pH 9.0; 10 mM Tris-Cl). To help reduce bias introduced by PCR, when constructing libraries for shotgun gut metagenome sequencing, libraries were prepared without PCR amplification. Libraries were sequenced at at Novogene (Nanjing, China) using the Illumina HiSeq (Illumina Inc., San Diego, California) yielding 150 base paired sequencing reads with data output of 6G per sample. Raw shotgun metagenomics sequence data was pre-processed through the MEDUSA pipeline.^77^ The metagenomic analysis yielded a mean of 23.4 ± 2.2 million reads per sample, with *16.6 ± 1.8 million of these reads successfully aligning*. 98% of sequencing reads met quality criteria, with only a *fraction (0.04%)* aligning with human genome. Gut metagenome sequences were processed by trimming and quality-checked with fastp v0.23.2 (setting ‘– detect_adapter_for_pe’).^78^ Following, we used Kraken v2.1.2,^79^ and PlusPF – 16 index (v20230605)^80^ for sequence analysis and taxonomic profiling of bacterial, viral, plasmid, fungal, archaeal, protozoal, and human components.

## Supplementary Results

### Patient and plasma biomarkers

**Figure S2.**
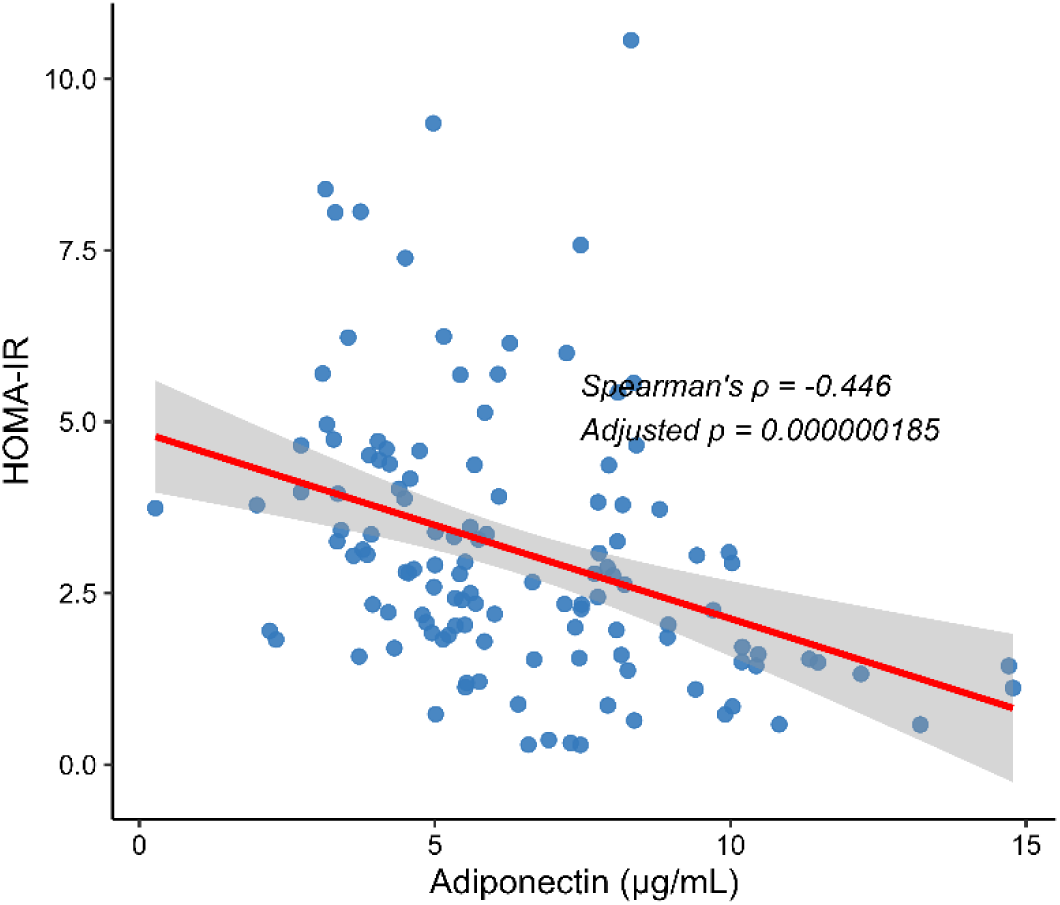
Correlation plots of adiponectin with HOMA-IR with lines of best fit and confidence interval. Spearman’s rank correlation (r), and its Benjamini-Hochberg adjusted p-value (*p*).

## Transcriptomics

**Figure S3.**
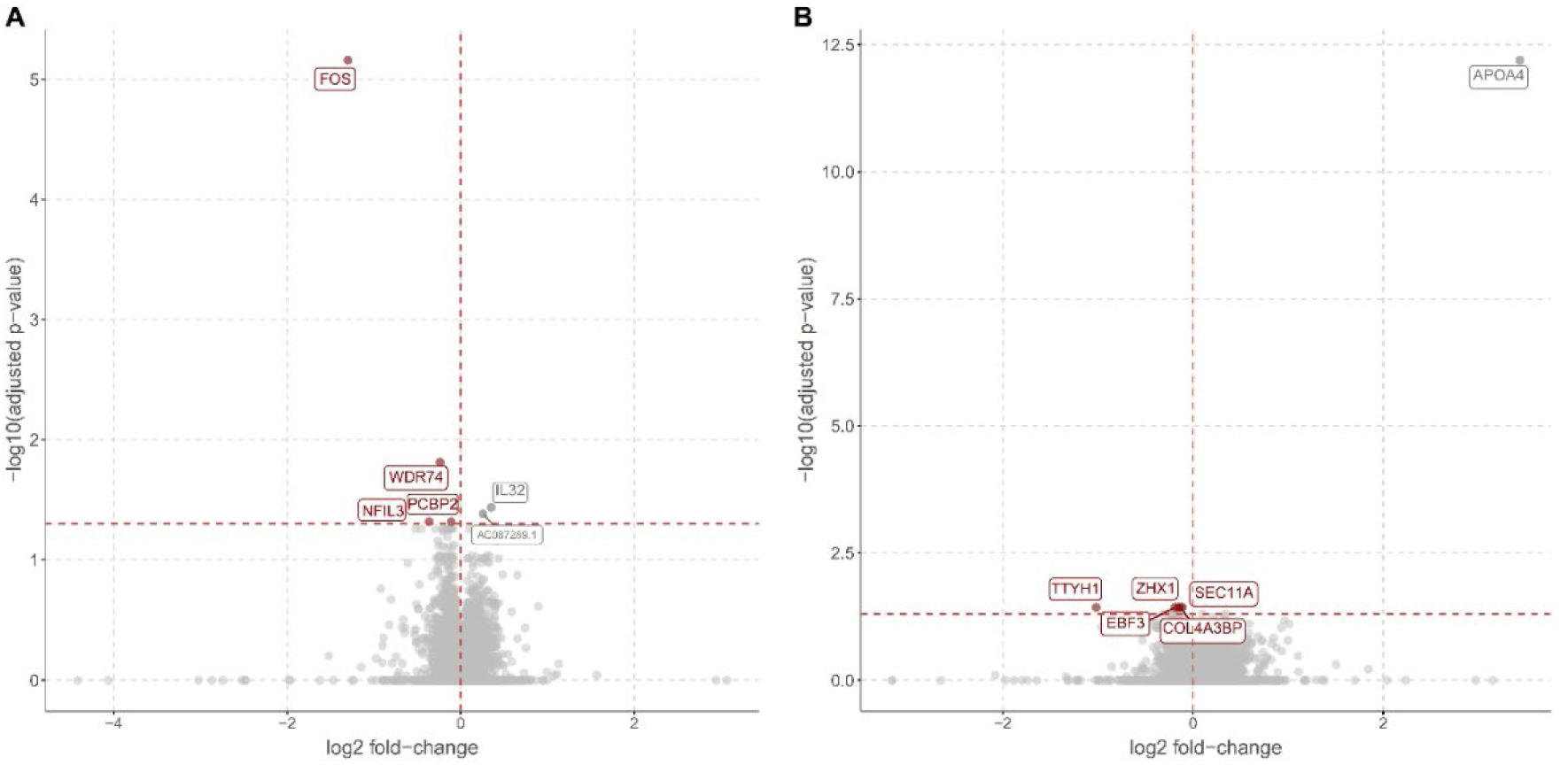
Volcano plots of adipose tissue transcriptomics in relation to dyslipidemia. A. Subcutaneous adipose tissue: genes on the left of center (red) downregulated in dyslipidemia, genes on the right (gray) upregulated. B. Visceral adipose tissue.

An overview of genes and their functional implication in lipid metabolism, (adipose tissue) inflammation, adipogenesis, and reactive oxygen species is seen in **Table S1** and **S2**.

**Table S1.**
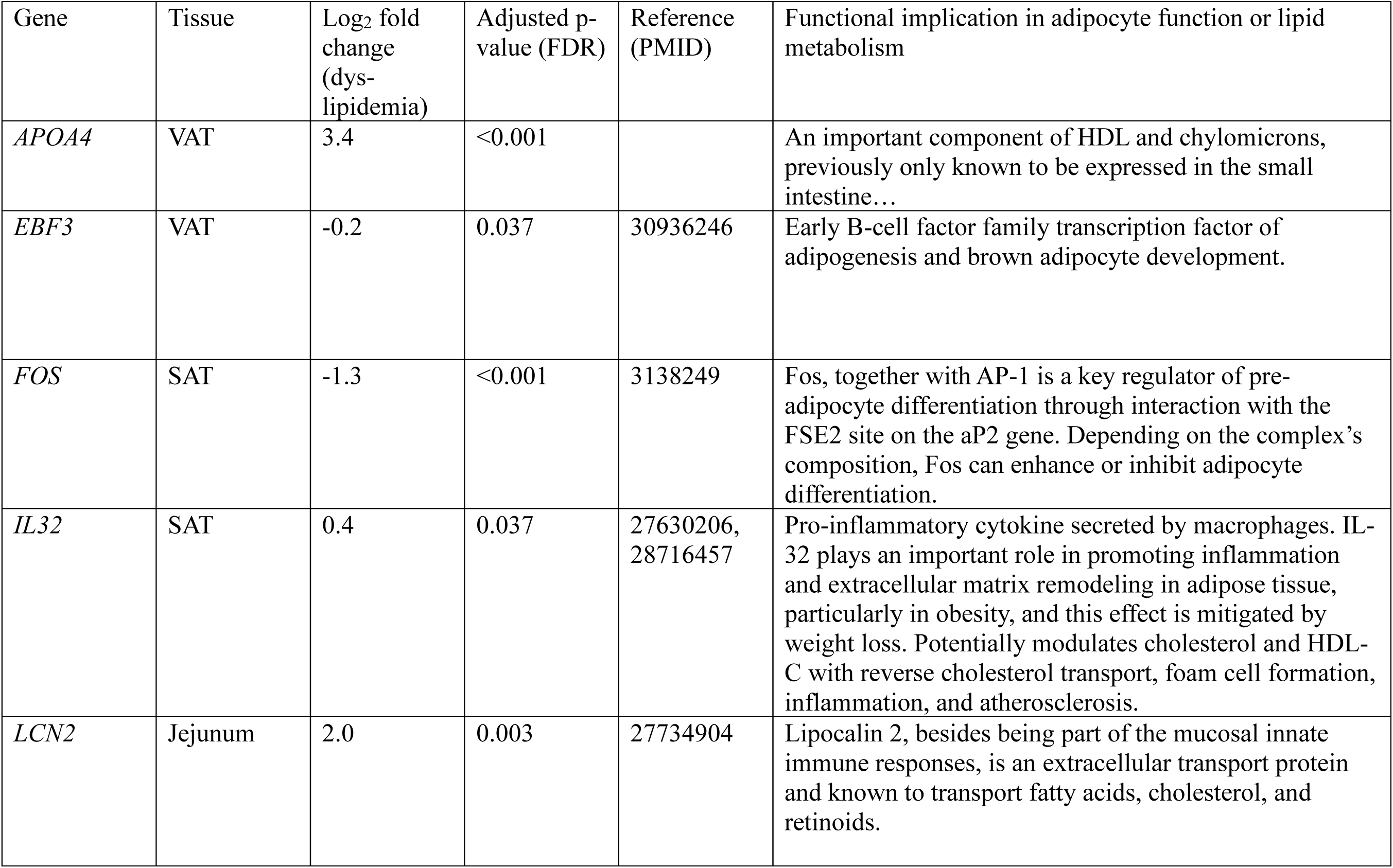
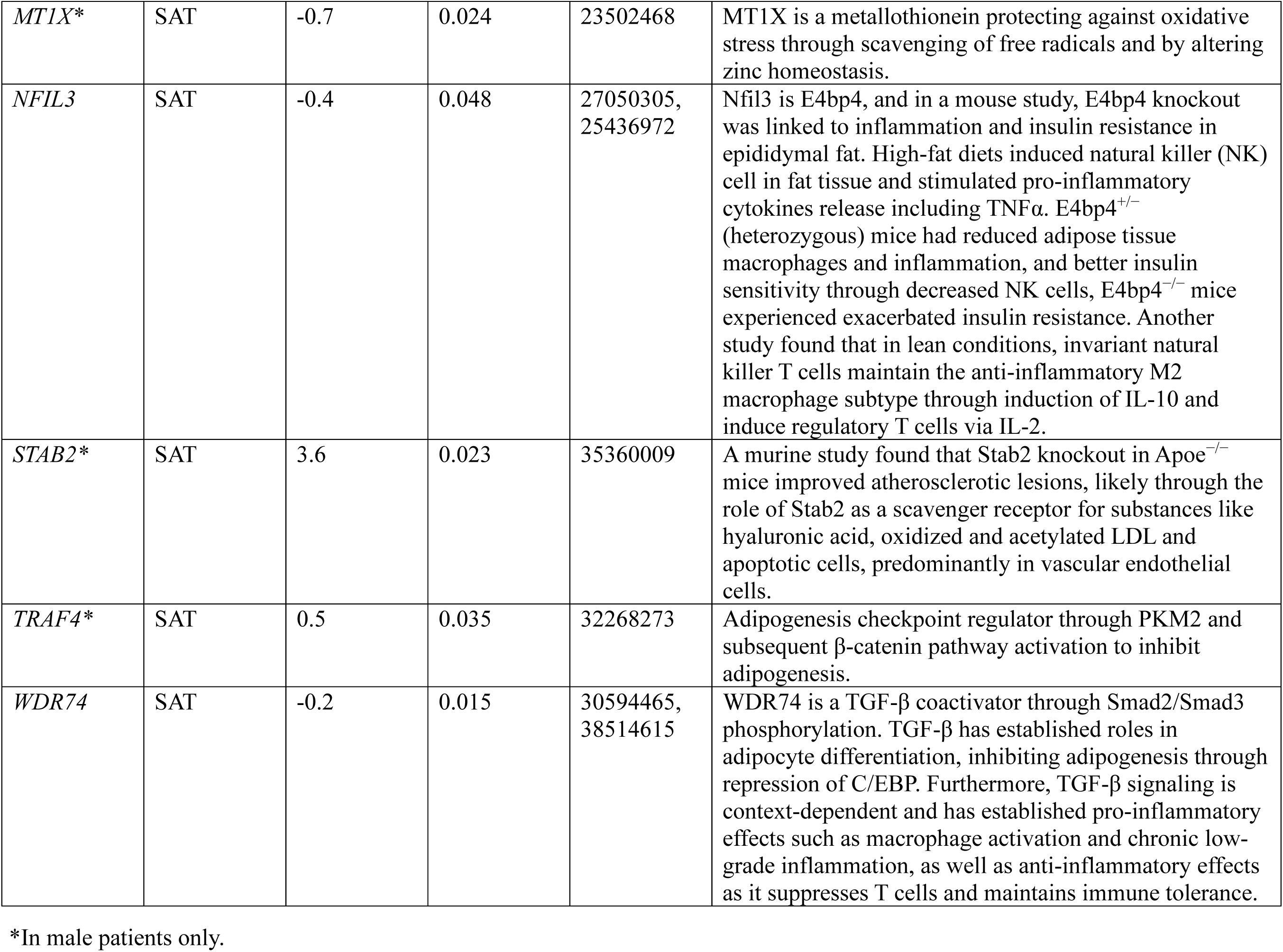
Differential gene expression across tissues in association with dyslipidemia (DESeq2 analysis, FDR-adjusted).

| Gene | Tissue | Log<sub>2</sub> fold change (dys-lipidemia) | Adjusted p-value (FDR) | Reference (PMID) | Functional implication in adipocyte function or lipid metabolism |
| --- | --- | --- | --- | --- | --- |
| *APOA4* | VAT | 3.4 | <0.001 |  | An important component of HDL and chylomicrons, previously only known to be expressed in the small intestine... |
| *EBF3* | VAT | -0.2 | 0.037 | 30936246 | Early B-cell factor family transcription factor of adipogenesis and brown adipocyte development. |
| *FOS* | SAT | -1.3 | <0.001 | 3138249 | Fos, together with AP-1 is a key regulator of pre-adipocyte differentiation through interaction with the FSE2 site on the aP2 gene. Depending on the complex's composition, Fos can enhance or inhibit adipocyte differentiation. |
| *IL32* | SAT | 0.4 | 0.037 | 27630206, 28716457 | Pro-inflammatory cytokine secreted by macrophages. IL-32 plays an important role in promoting inflammation and extracellular matrix remodeling in adipose tissue, particularly in obesity, and this effect is mitigated by weight loss. Potentially modulates cholesterol and HDL-C with reverse cholesterol transport, foam cell formation, inflammation, and atherosclerosis. |
| *LCN2* | Jejunum | 2.0 | 0.003 | 27734904 | Lipocalin 2, besides being part of the mucosal innate immune responses, is an extracellular transport protein and known to transport fatty acids, cholesterol, and retinoids. |
| *MT1X* * | SAT | -0.7 | 0.024 | 23502468 | MT1X is a metallothionein protecting against oxidative stress through scavenging of free radicals and by altering zinc homeostasis. |
| *NFIL3* | SAT | -0.4 | 0.048 | 27050305,<br>25436972 | Nfil3 is E4bp4, and in a mouse study, E4bp4 knockout was linked to inflammation and insulin resistance in epididymal fat. High-fat diets induced natural killer (NK) cell in fat tissue and stimulated pro-inflammatory cytokines release including TNFα. E4bp4<sup>+/-</sup> (heterozygous) mice had reduced adipose tissue macrophages and inflammation, and better insulin sensitivity through decreased NK cells, E4bp4<sup>-/-</sup> mice experienced exacerbated insulin resistance. Another study found that in lean conditions, invariant natural killer T cells maintain the anti-inflammatory M2 macrophage subtype through induction of IL-10 and induce regulatory T cells via IL-2. |
| *STAB2* * | SAT | 3.6 | 0.023 | 35360009 | A murine study found that Stab2 knockout in Apoe<sup>-/-</sup> mice improved atherosclerotic lesions, likely through the role of Stab2 as a scavenger receptor for substances like hyaluronic acid, oxidized and acetylated LDL and apoptotic cells, predominantly in vascular endothelial cells. |
| *TRAF4* * | SAT | 0.5 | 0.035 | 32268273 | Adipogenesis checkpoint regulator through PKM2 and subsequent β-catenin pathway activation to inhibit adipogenesis. |
| *WDR74* | SAT | -0.2 | 0.015 | 30594465,<br>38514615 | WDR74 is a TGF-β coactivator through Smad2/Smad3 phosphorylation. TGF-β has established roles in adipocyte differentiation, inhibiting adipogenesis through repression of C/EBP. Furthermore, TGF-β signaling is context-dependent and has established pro-inflammatory effects such as macrophage activation and chronic low-grade inflammation, as well as anti-inflammatory effects as it suppresses T cells and maintains immune tolerance. |

\*In male patients only.

**Table S2.**
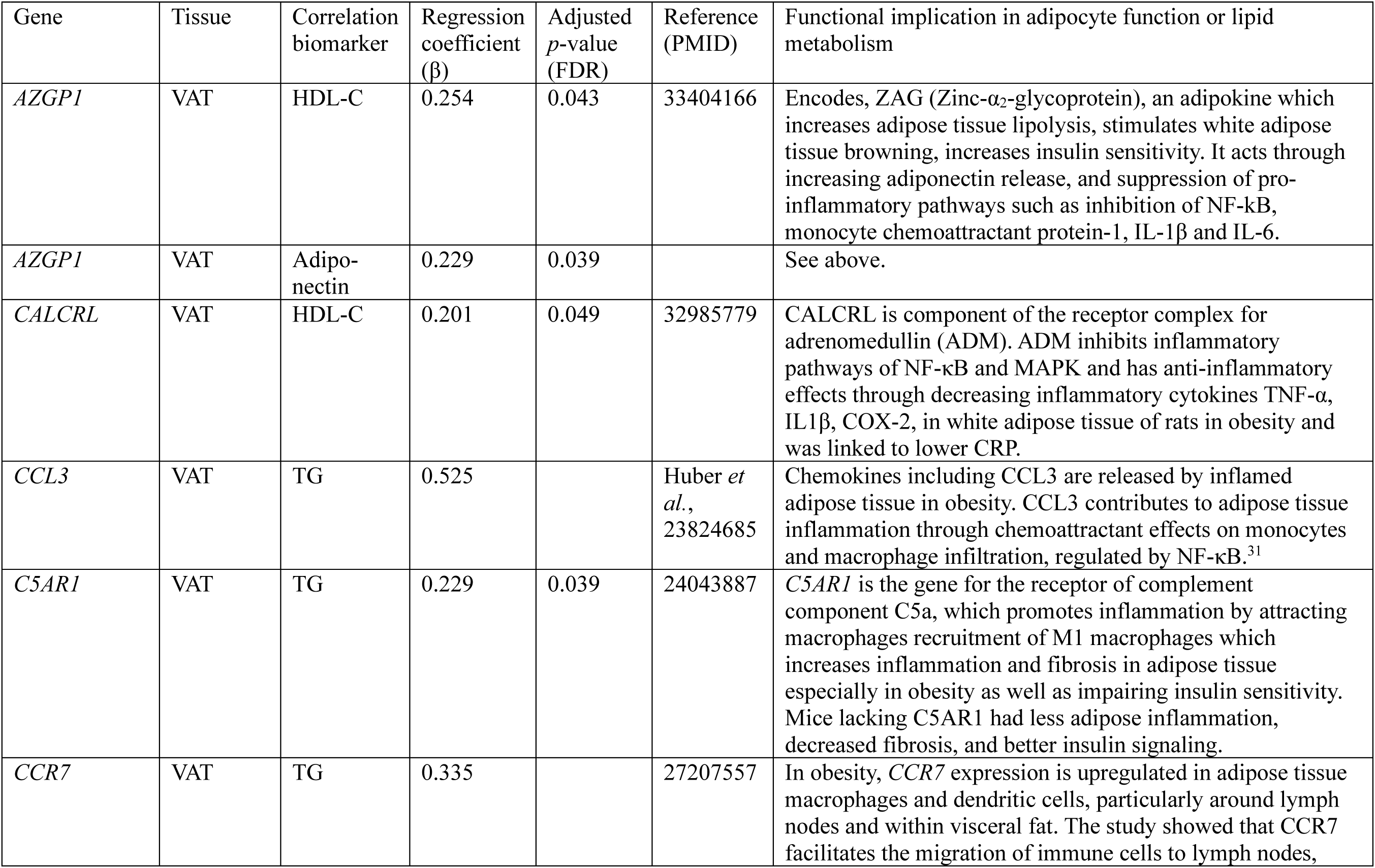
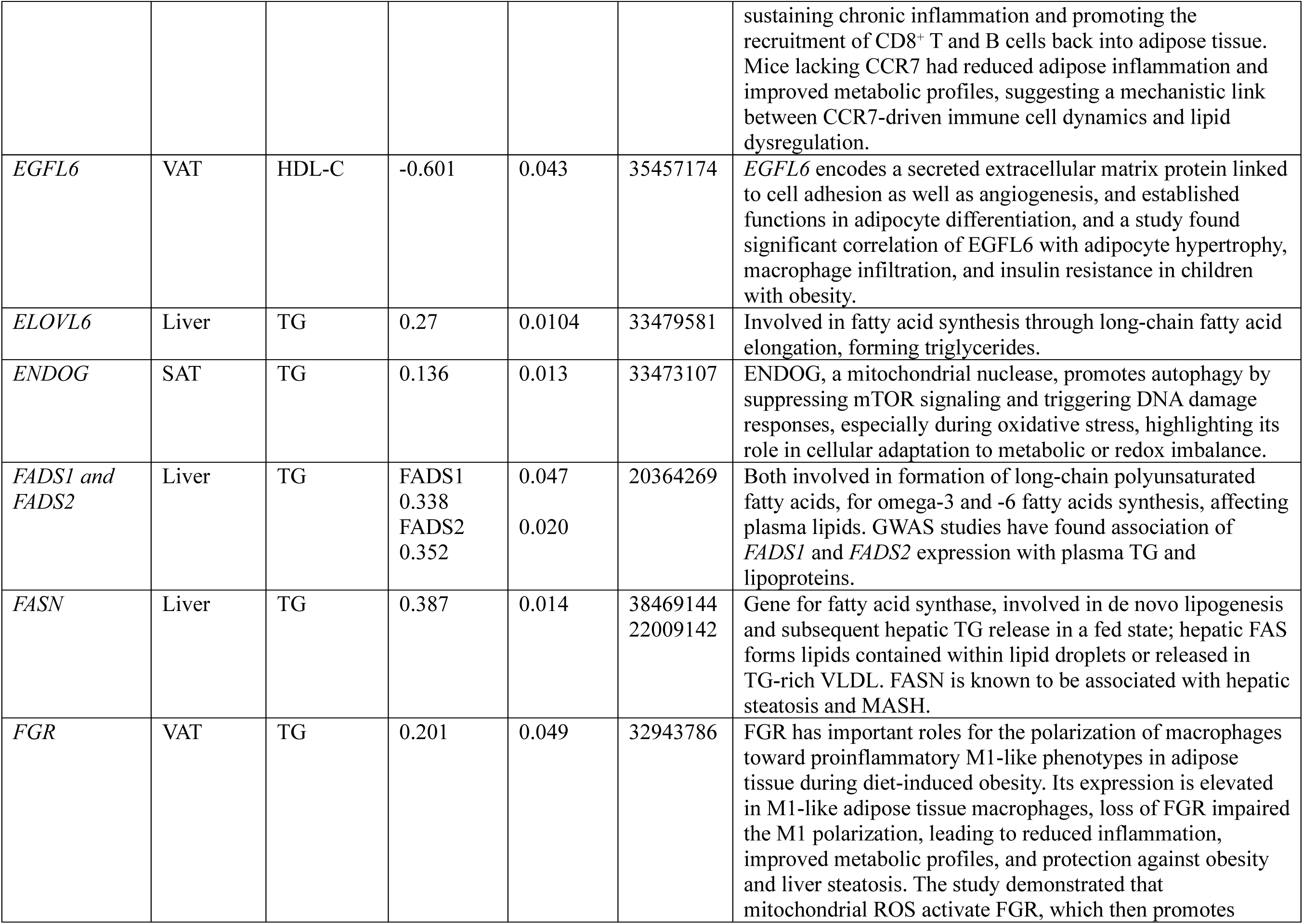
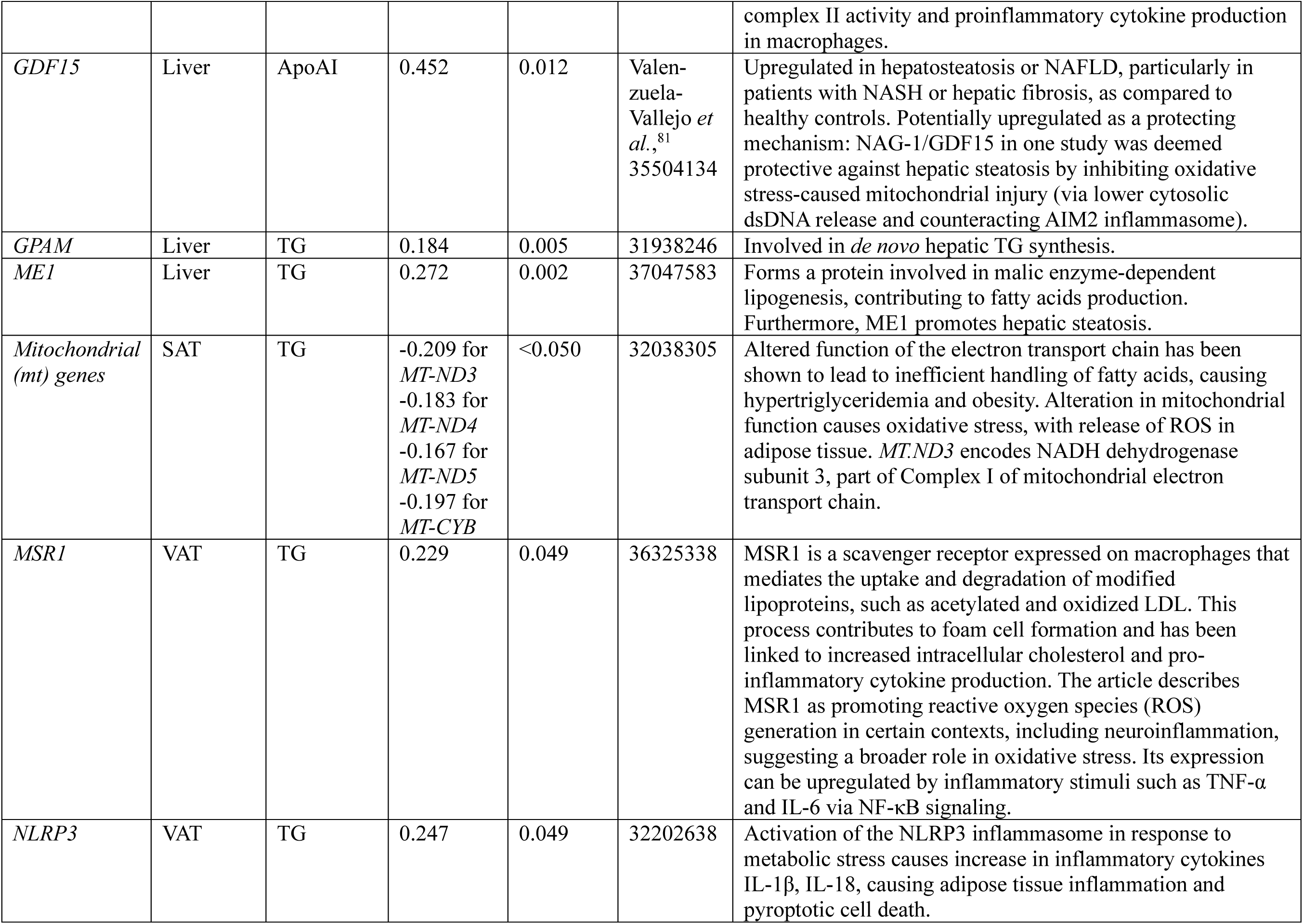
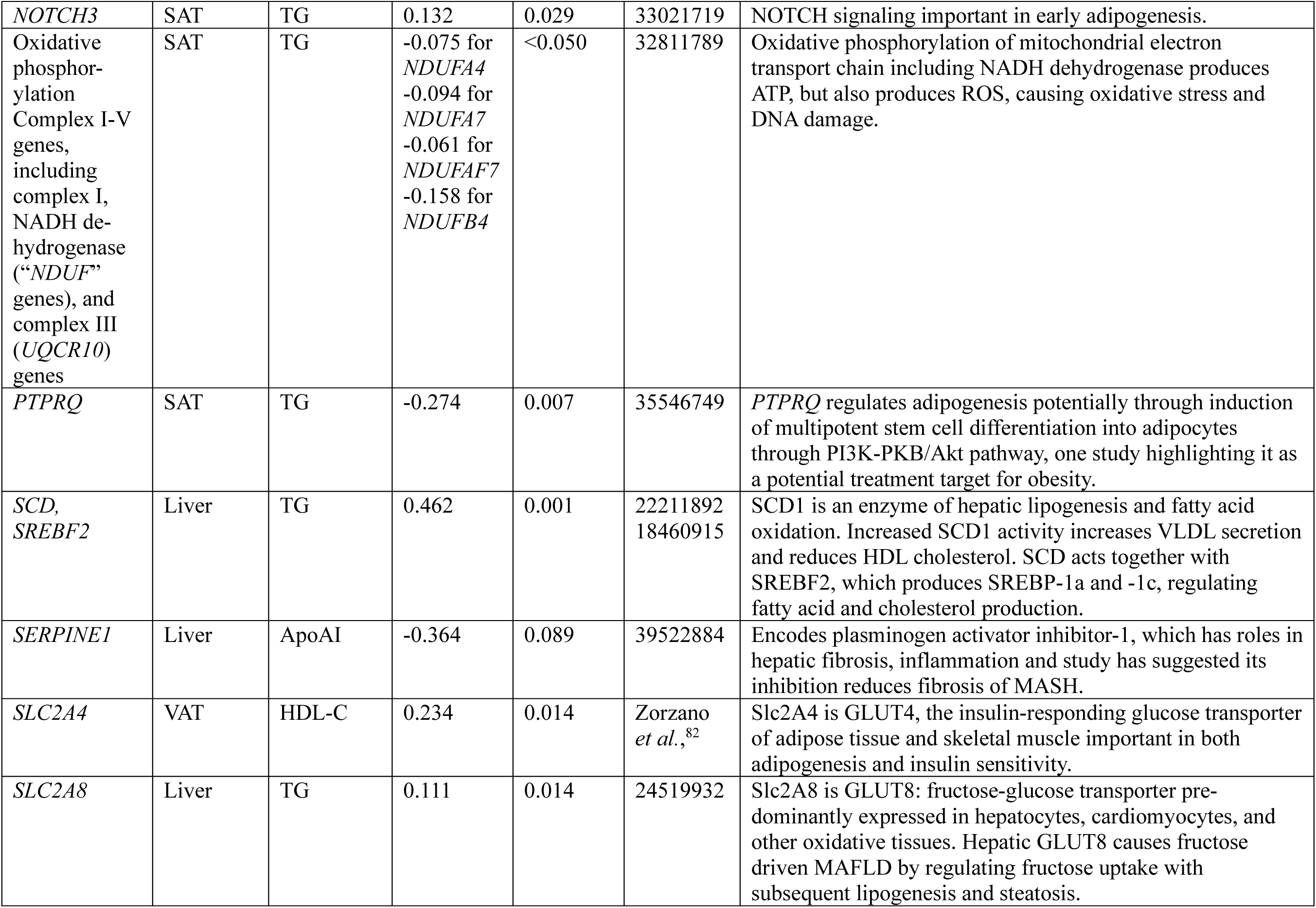
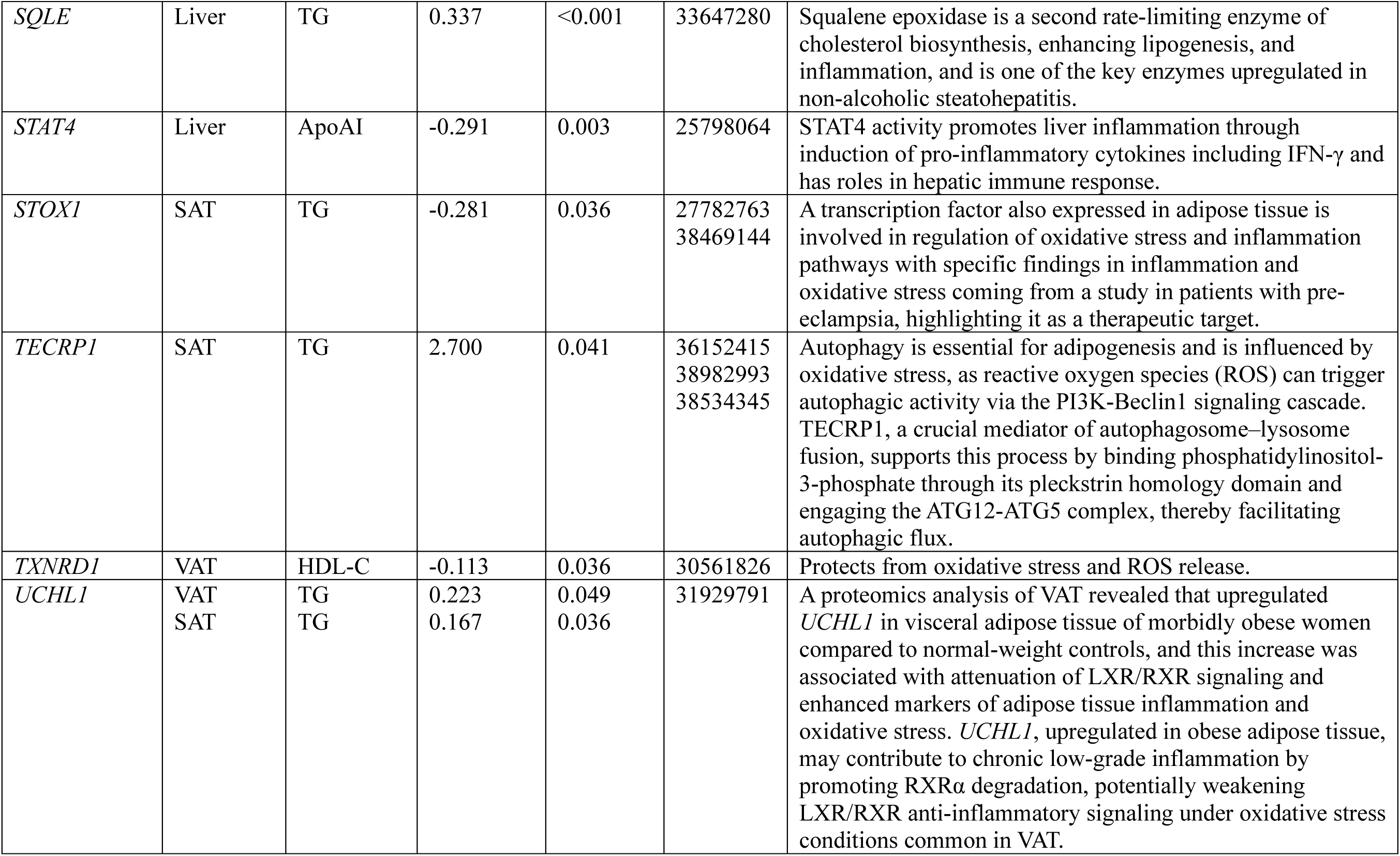
Gene expression associations with continuous lipids (MaAsLin linear modeling, FDR-adjusted).

**Figure S4.**
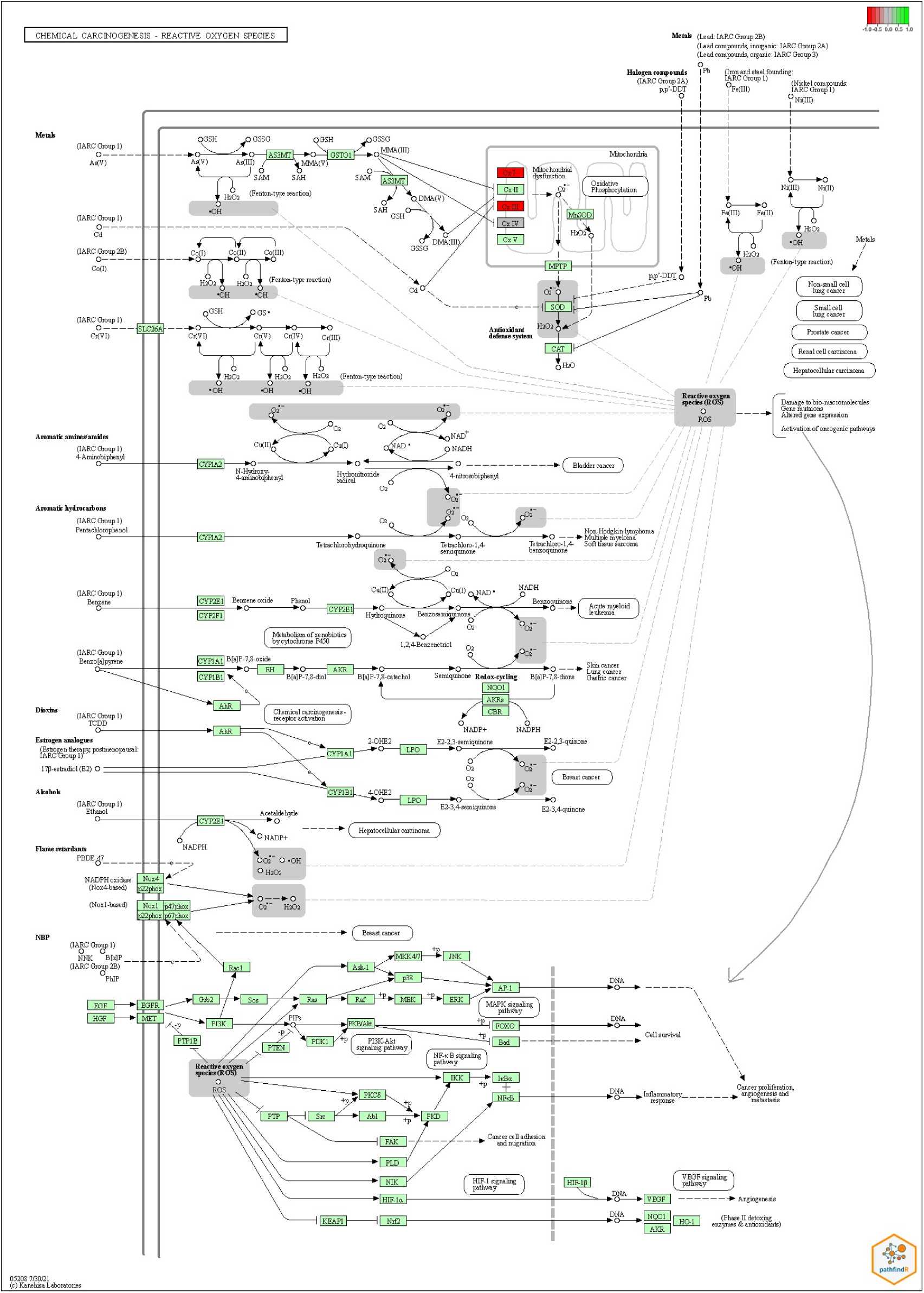
Chemical carcinogenesis-ROS production: Relative gene expression pattern of SAT genes of the chemical carcinogenesis-ROS production pathway in relation to plasma triglycerides. The color coding is from –1 (red) downregulated genes to +1 (green) upregulated genes where –1 and +1 signifies a log2 fold change.

## Metabolomics

**Figure S5.**
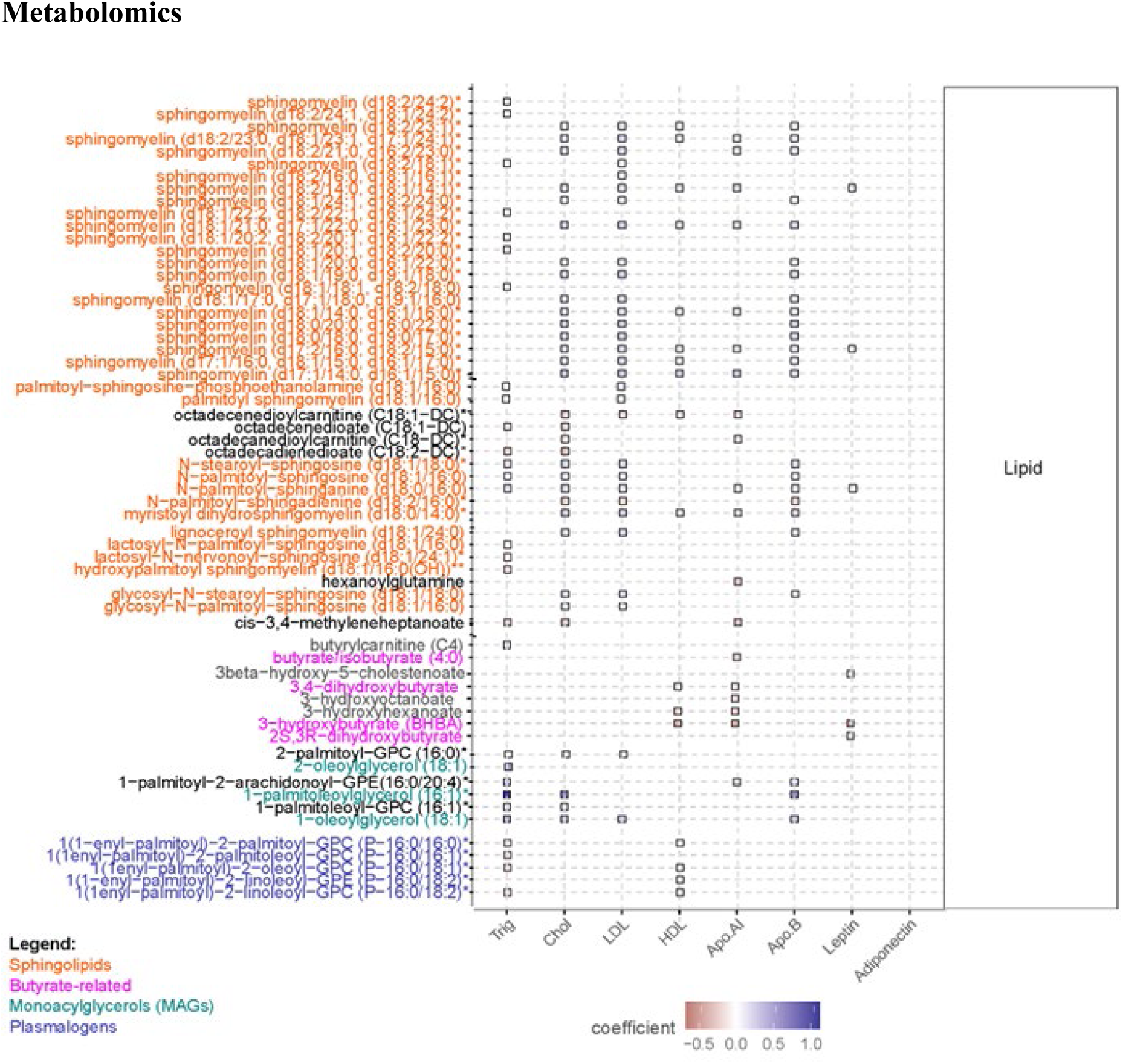
Selection of lipid metabolites in relation to plasma lipids, lipoproteins, and adipokines. The coefficient refers to the MaAsLin2 correlation. Only significant (p<0.05) associations depicted. Overview of all metabolites can be seen in Figure S6.

**Figure S6.**
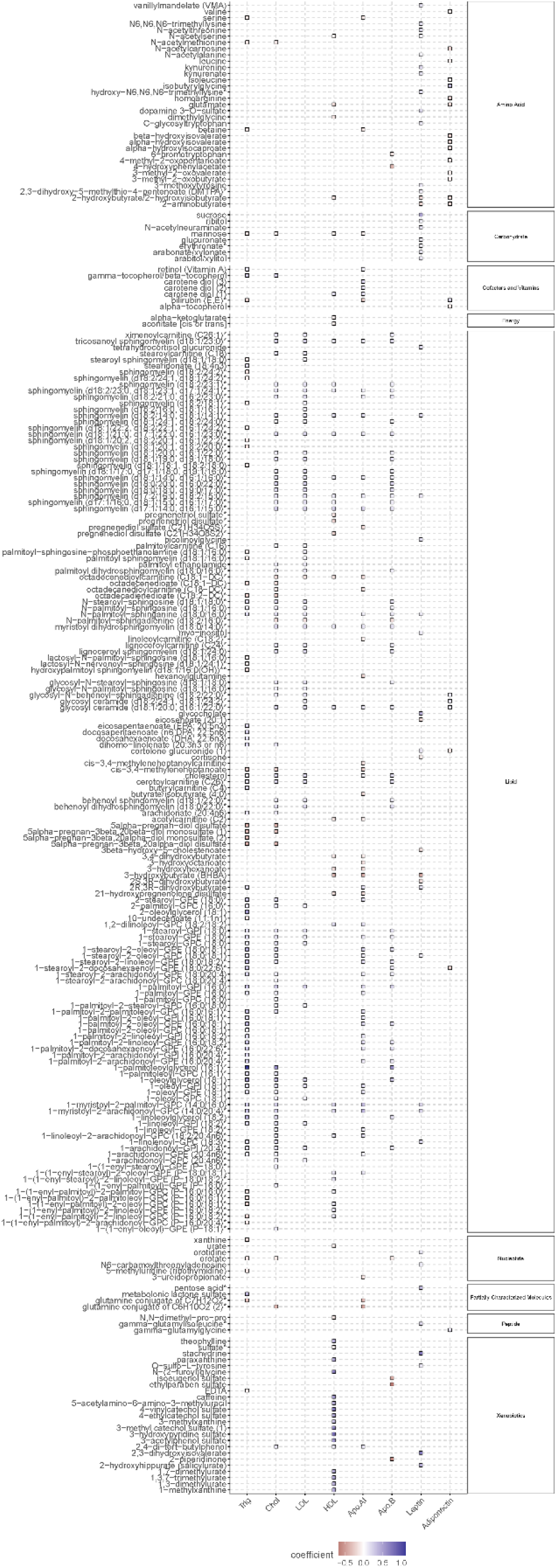
Metabolites in relation to plasma lipids, lipoproteins, and adipokines. The coefficient refers to the MaAsLin2 correlation. Only significant (*p*<0.05) associations depicted.

### Gut metagenomics

Principal coordinates analysis (PCoA) of gut microbiota (GM) found only modest difference in relative abundance of GM composition in patients with versus without dyslipidemia (*p* = 0.015, **Figure S7A**). Principal Component Analysis (PCA) however did not show difference in relative abundance between the groups (*p* = 0.156, **Figure S7B**). ANCOM-BC analysis found *Anaerococcus senegalensis* more abundant in dyslipidemia after adjusting for age and sex (*p* = 0.0125). DESeq2 analysis measuring relative abundance of bacterial species, highlighted species with significant change (**Figure S8**).

**Figure S7.**
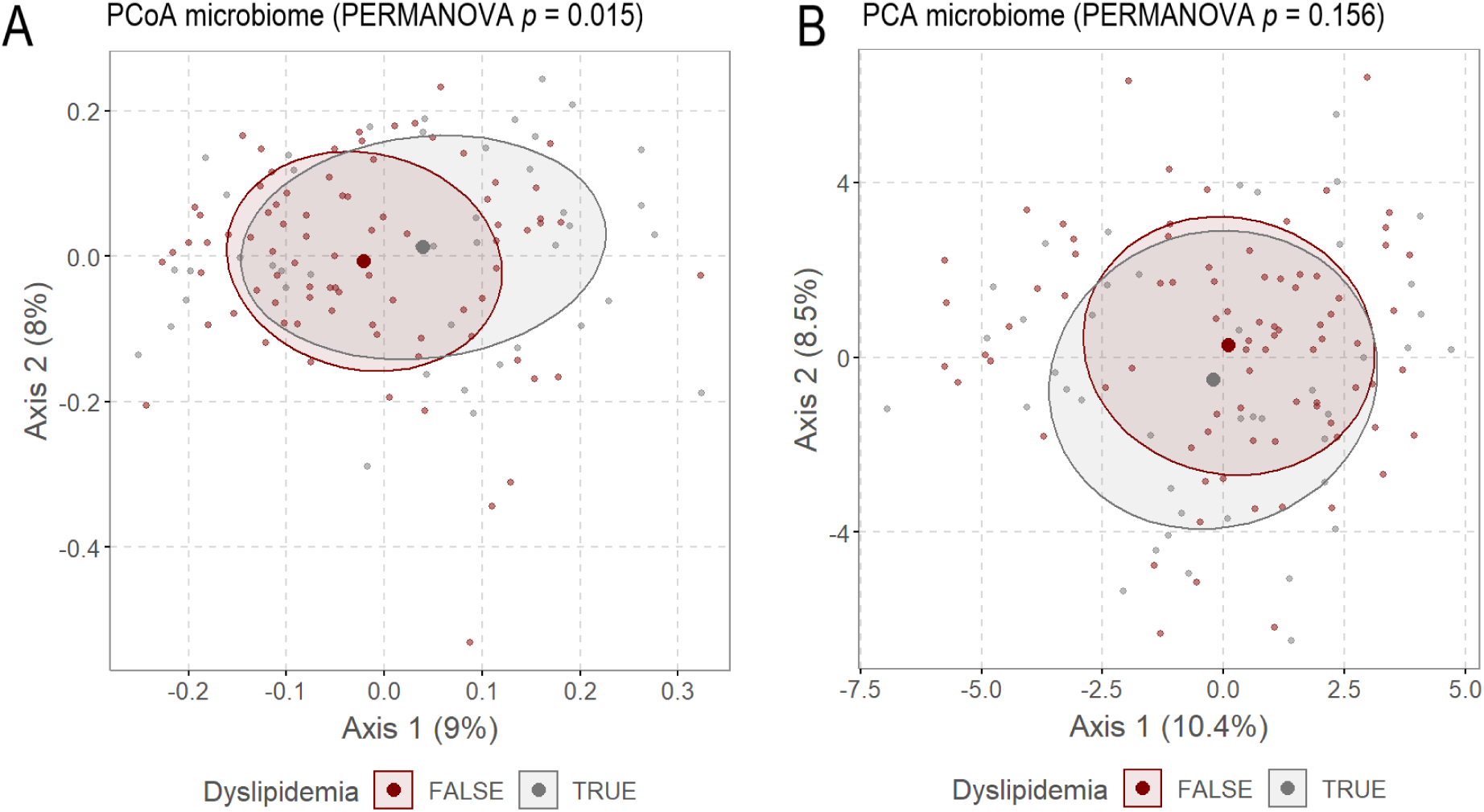
Difference in relative abundance of GM in patients with and without dyslipidemia. A. Principal Coordinates Analysis (PCoA) comparing relative abundance of GM composition between individuals with and without dyslipidemia. PCoA utilizes the Bray-Curtis distance matrix to illustrate compositional dissimilarity between the two groups, showing a modest difference of GM composition (PERMANOVA *p* = 0.015). B. Principal Component Analysis (PCA) focuses on variance across samples and utilizes centered log-ratio (clr) transformation to change relative abundances into a form analyzable by Euclidean geometry. PCA showed overlap without significant difference (PERMANOVA *p* = 0.156).

**Figure S8.**
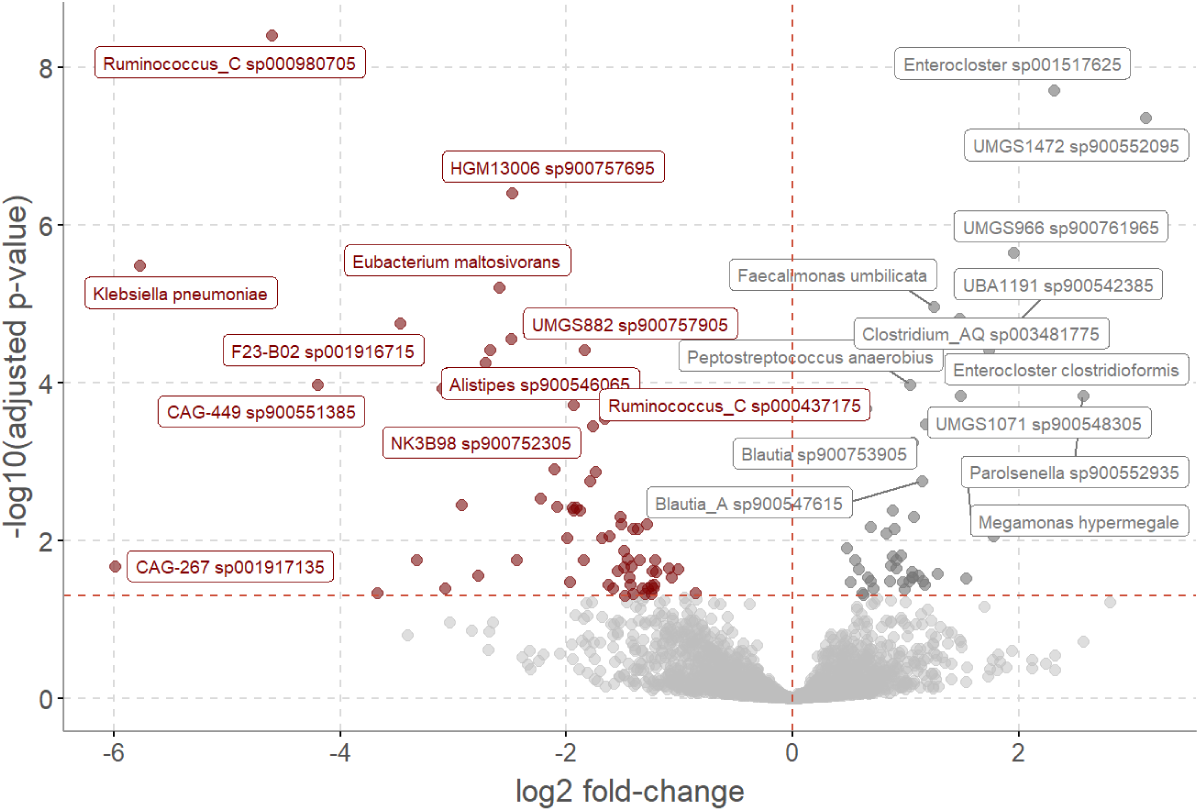
Volcano plot using DESeq2 to show the differential abundance of bacterial species associated with dyslipidemia, adjusted for age and sex. The x-axis represents log2 fold-change in abundance: species to the left have lower abundance in dyslipidemia, while those to the right have higher abundance. The y-axis indicates statistical significance (-log10 adjusted *p*-value). The y-axis shows the -log10 adjusted *p*-value, indicating the statistical significance of these changes.

**Figure S9.**
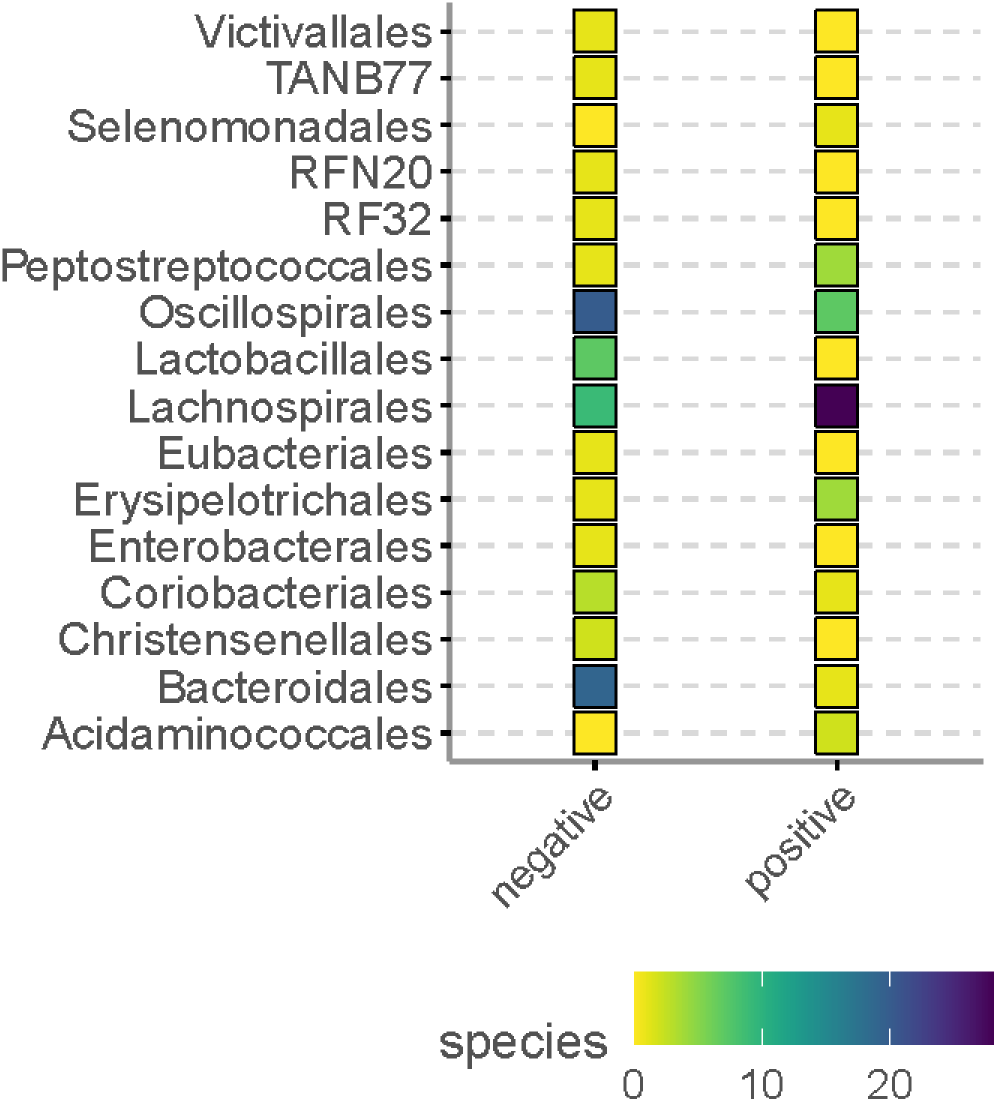
Differential abundance of bacterial orders from gut microbiota metagenomics in relation to dyslipidemia status. Illustrating number of species within each bacterial order in association with either the presence (positive) or absence (negative) of dyslipidemia.

## References

1. Blüher M. Obesity: global epidemiology and pathogenesis. Nat Rev Endocrinol. 2019;15(5):288–298. doi:10.1038/s41574-019-0176-8

2. GBD 2019 Risk Factors Collaborators. Global burden of 87 risk factors in 204 countries and territories, 1990-2019: a systematic analysis for the Global Burden of Disease Study 2019. Lancet Lond Engl. 2020;396(10258):1223–1249. doi:10.1016/S0140-6736(20)30752-2

3. Scheja L, Heeren J. The endocrine function of adipose tissues in health and cardiometabolic disease. Nat Rev Endocrinol. 2019;15(9):507–524. doi:10.1038/s41574-019-0230-6

4. Oikonomou EK, Antoniades C. The role of adipose tissue in cardiovascular health and disease. Nat Rev Cardiol. 2019;16(2):83–99. doi:10.1038/s41569-018-0097-6

5. Tacke F, Horn P, Wong VWS, et al. EASL–EASD–EASO Clinical Practice Guidelines on the management of metabolic dysfunction-associated steatotic liver disease (MASLD). J Hepatol. 2024;81(3):492–542. doi:10.1016/j.jhep.2024.04.031

6. Zwartjes MSZ, Gerdes VEA, Nieuwdorp M. The Role of Gut Microbiota and Its Produced Metabolites in Obesity, Dyslipidemia, Adipocyte Dysfunction, and Its Interventions. Metabolites. 2021;11(8):531. doi:10.3390/metabo11080531

7. Bays HE, Toth PP, Kris-Etherton PM, et al. Obesity, adiposity, and dyslipidemia: a consensus statement from the National Lipid Association. J Clin Lipidol. 2013;7(4):304–383. doi:10.1016/j.jacl.2013.04.001

8. Bays HE. “Sick Fat,” Metabolic Disease, and Atherosclerosis. Am J Med. 2009;122(1):S26–S37. doi:10.1016/j.amjmed.2008.10.015

9. Funcke JB, Scherer PE. Beyond adiponectin and leptin: adipose tissue-derived mediators of inter-organ communication. J Lipid Res. 2019;60(10):1648–1684. doi:10.1194/jlr.R094060

10. Tang L, Zhang F, Tong N. The association of visceral adipose tissue and subcutaneous adipose tissue with metabolic risk factors in a large population of Chinese adults. Clin Endocrinol (Oxf). 2016;85(1):46–53. doi:10.1111/cen.13013

11. Fox CS, Massaro JM, Hoffmann U, et al. Abdominal visceral and subcutaneous adipose tissue compartments: association with metabolic risk factors in the Framingham Heart Study. Circulation. 2007;116(1):39–48. doi:10.1161/CIRCULATIONAHA.106.675355

12. von Krüchten R, Lorbeer R, Müller-Peltzer K, et al. Association between Adipose Tissue Depots and Dyslipidemia: The KORA-MRI Population-Based Study. Nutrients. 2022;14(4):797. doi:10.3390/nu14040797

13. Xu S, Lu F, Gao J, Yuan Y. Inflammation-mediated metabolic regulation in adipose tissue. Obes Rev Off J Int Assoc Study Obes. 2024;25(6):e13724. doi:10.1111/obr.13724

14. Landecho MF, Tuero C, Valentí V, Bilbao I, de la Higuera M, Frühbeck G. Relevance of Leptin and Other Adipokines in Obesity-Associated Cardiovascular Risk. Nutrients. 2019;11(11):2664. doi:10.3390/nu11112664

15. Franssens BT, Westerink J, van der Graaf Y, Nathoe HM, Visseren FLJ, SMART study group. Metabolic consequences of adipose tissue dysfunction and not adiposity per se increase the risk of cardiovascular events and mortality in patients with type 2 diabetes. Int J Cardiol. 2016;222:72–77. doi:10.1016/j.ijcard.2016.07.081

16. Liu L, Shi Z, Ji X, et al. Adipokines, adiposity, and atherosclerosis. Cell Mol Life Sci CMLS. 2022;79(5):272. doi:10.1007/s00018-022-04286-2

17. Chaurasiya V, Nidhina Haridas PA, Olkkonen VM. Adipocyte-endothelial cell interplay in adipose tissue physiology. Biochem Pharmacol. 2024;222:116081. doi:10.1016/j.bcp.2024.116081

18. Kurilshikov A, van den Munckhof ICL, Chen L, et al. Gut Microbial Associations to Plasma Metabolites Linked to Cardiovascular Phenotypes and Risk. Circ Res. 2019;124(12):1808–1820. doi:10.1161/CIRCRESAHA.118.314642

19. Van Olden CC, Van de Laar AW, Meijnikman AS, et al. A systems biology approach to understand gut microbiota and host metabolism in morbid obesity: design of the BARIA Longitudinal Cohort Study. J Intern Med. 2021;289(3):340–354. doi:10.1111/joim.13157

20. Expert Panel on Detection, Evaluation, and Treatment of High Blood Cholesterol in Adults. Executive Summary of The Third Report of The National Cholesterol Education Program (NCEP) Expert Panel on Detection, Evaluation, And Treatment of High Blood Cholesterol In Adults (Adult Treatment Panel III). JAMA. 2001;285(19):2486–2497. doi:10.1001/jama.285.19.2486

21. Friedewald WT, Levy RI, Fredrickson DS. Estimation of the concentration of low-density lipoprotein cholesterol in plasma, without use of the preparative ultracentrifuge. Clin Chem. 1972;18(6):499–502.

22. Ulgen E, Ozisik O, Sezerman OU. pathfindR: An R Package for Comprehensive Identification of Enriched Pathways in Omics Data Through Active Subnetworks. Front Genet. 2019;10:858. doi:10.3389/fgene.2019.00858

23. The Comprehensive R Archive Network. Accessed May 5, 2023. https://cran.r-project.org/

24. Posit. RStudio Desktop: RStudio-2024.04.2-764. Published online June 2024. Accessed August 16, 2024. https://posit.co/download/rstudio-desktop/

25. Klop B, Elte JWF, Cabezas MC. Dyslipidemia in obesity: mechanisms and potential targets. Nutrients. 2013;5(4):1218–1240. doi:10.3390/nu5041218

26. Deng Z, Fan T, Xiao C, et al. TGF-β signaling in health, disease and therapeutics. Signal Transduct Target Ther. 2024;9(1):1–40. doi:10.1038/s41392-024-01764-w

27. Lee BC, Kim MS, Pae M, et al. Adipose Natural Killer Cells Regulate Adipose Tissue Macrophages to Promote Insulin Resistance in Obesity. Cell Metab. 2016;23(4):685–698. doi:10.1016/j.cmet.2016.03.002

28. Lynch L, Michelet X, Zhang S, et al. Regulatory iNKT cells lack expression of the transcription factor PLZF and control the homeostasis of T(reg) cells and macrophages in adipose tissue. Nat Immunol. 2015;16(1):85–95. doi:10.1038/ni.3047

29. Kayashima Y, Clanton CA, Lewis AM, et al. Reduction of Stabilin-2 Contributes to a Protection Against Atherosclerosis. Front Cardiovasc Med. 2022;9:818662. doi:10.3389/fcvm.2022.818662

30. Shang C, Sun W, Wang C, et al. Comparative Proteomic Analysis of Visceral Adipose Tissue in Morbidly Obese and Normal Weight Chinese Women. Int J Endocrinol. 2019;2019:2302753. doi:10.1155/2019/2302753

31. Huber J, Kiefer F, Zeyda M, et al. CCL2 and CCL3 chemokine gene expression and their receptors in human visceral and subcutaneous adipose tissue is associated with obesity and insulin resistance. Exp Clin Endocrinol Diabetes. 2007;115(S 1):P02_063. doi:10.1055/s-2007-972470

32. Tourniaire F, Romier-Crouzet B, Lee JH, et al. Chemokine Expression in Inflamed Adipose Tissue Is Mainly Mediated by NF-κB. PLoS ONE. 2013;8(6):e66515. doi:10.1371/journal.pone.0066515

33. Griffin MJ. On the Immunometabolic Role of NF-κB in Adipocytes. Immunometabolism. 2022;4(1):e220003. doi:10.20900/immunometab20220003

34. Namkhah Z, Naeini F, Ostadrahimi A, Tutunchi H, Hosseinzadeh-Attar MJ. The association of the adipokine zinc-alpha2-glycoprotein with non-alcoholic fatty liver disease and related risk factors: A comprehensive systematic review. Int J Clin Pract. 2021;75(7):e13985. doi:10.1111/ijcp.13985

35. Hellmann J, Sansbury BE, Holden CR, et al. CCR7 Maintains Nonresolving Lymph Node and Adipose Inflammation in Obesity. Diabetes. 2016;65(8):2268–2281. doi:10.2337/db15-1689

36. Phieler J, Chung KJ, Chatzigeorgiou A, et al. The complement anaphylatoxin C5a receptor contributes to obese adipose tissue inflammation and insulin resistance. J Immunol Baltim Md 1950. 2013;191(8):4367–4374. doi:10.4049/jimmunol.1300038

37. Barra NG, Henriksbo BD, Anhê FF, Schertzer JD. The NLRP3 inflammasome regulates adipose tissue metabolism. Biochem J. 2020;477(6):1089–1107. doi:10.1042/BCJ20190472

38. Spiegelman BM, Distel RJ, Ro HS, Rosen BS, Satterberg B. fos protooncogene and the regulation of gene expression in adipocyte differentiation. J Cell Biol. 1988;107(3):829–832.

39. Liu MC, Logan H, Newman JJ. Distinct roles for Notch1 and Notch3 in human adipose-derived stem/stromal cell adipogenesis. Mol Biol Rep. 2020;47(11):8439–8450. doi:10.1007/s11033-020-05884-8

40. Zhang W, Tang Z, Fan S, et al. Protein Tyrosine Phosphatase Receptor-type Q: Structure, Activity, and Implications in Human Disease. Protein Pept Lett. 2022;29(7):567–573. doi:10.2174/0929866529666220511141826

41. Cen S, Li J, Cai Z, et al. TRAF4 acts as a fate checkpoint to regulate the adipogenic differentiation of MSCs by activating PKM2. EBioMedicine. 2020;54:102722. doi:10.1016/j.ebiom.2020.102722

42. Landgraf K, Kühnapfel A, Schlanstein M, et al. Transcriptome Analyses of Adipose Tissue Samples Identify EGFL6 as a Candidate Gene Involved in Obesity-Related Adipose Tissue Dysfunction in Children. Int J Mol Sci. 2022;23(8):4349. doi:10.3390/ijms23084349

43. Nono Nankam PA, Nguelefack TB, Goedecke JH, Blüher M. Contribution of Adipose Tissue Oxidative Stress to Obesity-Associated Diabetes Risk and Ethnic Differences: Focus on Women of African Ancestry. Antioxidants. 2021;10(4):622. doi:10.3390/antiox10040622

44. Khan F, Khan H, Khan A, et al. Autophagy in adipogenesis: Molecular mechanisms and regulation by bioactive compounds. Biomed Pharmacother. 2022;155:113715. doi:10.1016/j.biopha.2022.113715

45. Hong C, Li X, Zhang K, et al. Novel perspectives on autophagy-oxidative stress-inflammation axis in the orchestration of adipogenesis. Front Endocrinol. 2024;15:1404697. doi:10.3389/fendo.2024.1404697

46. Wang W, Li J, Tan J, et al. Endonuclease G promotes autophagy by suppressing mTOR signaling and activating the DNA damage response. Nat Commun. 2021;12:476. doi:10.1038/s41467-020-20780-2

47. Ruttkay-Nedecky B, Nejdl L, Gumulec J, et al. The Role of Metallothionein in Oxidative Stress. Int J Mol Sci. 2013;14(3):6044–6066. doi:10.3390/ijms14036044

48. Acín-Pérez R, Iborra S, Martí-Mateos Y, et al. Fgr kinase is required for proinflammatory macrophage activation during diet-induced obesity. Nat Metab. 2020;2(9):974–988. doi:10.1038/s42255-020-00273-8

49. Loboda A, Jozkowicz A, Dulak J. HIF-1 and HIF-2 transcription factors--similar but not identical. Mol Cells. 2010;29(5):435–442. doi:10.1007/s10059-010-0067-2

50. He Y, Luo Y, Zhang D, et al. PGK1-mediated cancer progression and drug resistance. Am J Cancer Res. 2019;9(11):2280–2302.

51. Ullah K, Ai L, Humayun Z, Wu R. Targeting Endothelial HIF2α/ARNT Expression for Ischemic Heart Disease Therapy. Biology. 2023;12(7):995. doi:10.3390/biology12070995

52. Lee D, Xu IMJ, Chiu DKC, et al. Induction of Oxidative Stress Through Inhibition of Thioredoxin Reductase 1 Is an Effective Therapeutic Approach for Hepatocellular Carcinoma. Hepatol Baltim Md. 2019;69(4):1768–1786. doi:10.1002/hep.30467

53. Liu D, Wong CC, Zhou Y, et al. Squalene Epoxidase Induces Nonalcoholic Steatohepatitis Via Binding to Carbonic Anhydrase III and is a Therapeutic Target. Gastroenterology. 2021;160(7):2467–2482.e3. doi:10.1053/j.gastro.2021.02.051

54. DeBosch BJ, Chen Z, Saben JL, Finck BN, Moley KH. Glucose transporter 8 (GLUT8) mediates fructose-induced de novo lipogenesis and macrosteatosis. J Biol Chem. 2014;289(16):10989–10998. doi:10.1074/jbc.M113.527002

55. Guillamat-Prats R, Rami M, Herzig S, Steffens S. Endocannabinoid Signalling in Atherosclerosis and Related Metabolic Complications. Thromb Haemost. 2019;119(4):567–575. doi:10.1055/s-0039-1678738

56. Lowe H, Toyang N, Steele B, Bryant J, Ngwa W. The Endocannabinoid System: A Potential Target for the Treatment of Various Diseases. Int J Mol Sci. 2021;22(17):9472. doi:10.3390/ijms22179472

57. Gaetani S, Kaye WH, Cuomo V, Piomelli D. Role of endocannabinoids and their analogues in obesity and eating disorders. Eat Weight Disord EWD. 2008;13(3):e42–48.

58. De Filippo C, Costa A, Becagli MV, Monroy MM, Provensi G, Passani MB. Gut microbiota and oleoylethanolamide in the regulation of intestinal homeostasis. Front Endocrinol. 2023;14. Accessed June 8, 2023. https://www.frontiersin.org/articles/10.3389/fendo.2023.1135157

59. Russo R, Cristiano C, Avagliano C, et al. Gut-brain Axis: Role of Lipids in the Regulation of Inflammation, Pain and CNS Diseases. Curr Med Chem. 2018;25(32):3930–3952. doi:10.2174/0929867324666170216113756

60. Rinne P, Guillamat-Prats R, Rami M, et al. Palmitoylethanolamide Promotes a Proresolving Macrophage Phenotype and Attenuates Atherosclerotic Plaque Formation. Arterioscler Thromb Vasc Biol. 2018;38(11):2562–2575. doi:10.1161/ATVBAHA.118.311185

61. Clayton P, Hill M, Bogoda N, Subah S, Venkatesh R. Palmitoylethanolamide: A Natural Compound for Health Management. Int J Mol Sci. 2021;22(10):5305. doi:10.3390/ijms22105305

62. Louca P, Meijnikman AS, Nogal A, et al. The secondary bile acid isoursodeoxycholate correlates with post-prandial lipemia, inflammation, and appetite and changes post-bariatric surgery. Cell Rep Med. 2023;4(4):100993. doi:10.1016/j.xcrm.2023.100993

63. Green CD, Maceyka M, Cowart LA, Spiegel S. Sphingolipids in metabolic disease: The good, the bad, and the unknown. Cell Metab. 2021;33(7):1293–1306. doi:10.1016/j.cmet.2021.06.006

64. Ohira H, Tsutsui W, Fujioka Y. Are Short Chain Fatty Acids in Gut Microbiota Defensive Players for Inflammation and Atherosclerosis? J Atheroscler Thromb. 2017;24(7):660–672. doi:10.5551/jat.RV17006

65. Amiri P, Hosseini SA, Ghaffari S, et al. Role of Butyrate, a Gut Microbiota Derived Metabolite, in Cardiovascular Diseases: A comprehensive narrative review. Front Pharmacol. 2022;12:837509. doi:10.3389/fphar.2021.837509

66. Paul S, Lancaster GI, Meikle PJ. Plasmalogens: A potential therapeutic target for neurodegenerative and cardiometabolic disease. Prog Lipid Res. 2019;74:186–195. doi:10.1016/j.plipres.2019.04.003

67. Park H, He A, Lodhi IJ. Lipid Regulators of Thermogenic Fat Activation. Trends Endocrinol Metab TEM. 2019;30(10):710–723. doi:10.1016/j.tem.2019.07.020

68. Warmbrunn MV, Boulund U, Aron-Wisnewsky J, et al. Networks of gut bacteria relate to cardiovascular disease in a multi-ethnic population: the HELIUS study. Cardiovasc Res. 2024;120(4):372–384. doi:10.1093/cvr/cvae018

69. Matthews DR, Hosker JP, Rudenski AS, Naylor BA, Treacher DF, Turner RC. Homeostasis model assessment: insulin resistance and beta-cell function from fasting plasma glucose and insulin concentrations in man. Diabetologia. 1985;28(7):412–419. doi:10.1007/BF00280883

70. Levy JC, Matthews DR, Hermans MP. Correct homeostasis model assessment (HOMA) evaluation uses the computer program. Diabetes Care. 1998;21(12):2191–2192. doi:10.2337/diacare.21.12.2191

71. Radcliffe Department of Medicine - Medical Sciences Division, University of Oxford. HOMA2 Calculator software. 2023. Accessed May 10, 2023. https://www.rdm.ox.ac.uk/about/our-clinical-facilities-and-mrc-units/DTU/software/homa/download

72. Bolger AM, Lohse M, Usadel B. Trimmomatic: a flexible trimmer for Illumina sequence data. Bioinformatics. 2014;30(15):2114–2120. doi:10.1093/bioinformatics/btu170

73. Near-optimal probabilistic RNA-seq quantification | Nature Biotechnology. Accessed August 9, 2024. https://www.nature.com/articles/nbt.3519

74. 74. Moderated estimation of fold change and dispersion for RNA-seq data with DESeq2 | Genome Biology | Full Text. Accessed August 9, 2024. https://genomebiology.biomedcentral.com/articles/10.1186/s13059-014-0550-8

75. Mallick H, Rahnavard A, McIver LJ, et al. Multivariable association discovery in population-scale meta-omics studies. PLoS Comput Biol. 2021;17(11):e1009442. doi:10.1371/journal.pcbi.1009442

76. Costea PI, Zeller G, Sunagawa S, et al. Towards standards for human fecal sample processing in metagenomic studies. Nat Biotechnol. 2017;35(11):1069–1076. doi:10.1038/nbt.3960

77. Morais DAA, Cavalcante JVF, Monteiro SS, Pasquali MAB, Dalmolin RJS. MEDUSA: A Pipeline for Sensitive Taxonomic Classification and Flexible Functional Annotation of Metagenomic Shotgun Sequences. Front Genet. 2022;13:814437. doi:10.3389/fgene.2022.814437

78. Chen S, Zhou Y, Chen Y, Gu J. fastp: an ultra-fast all-in-one FASTQ preprocessor. Bioinforma Oxf Engl. 2018;34(17):i884–i890. doi:10.1093/bioinformatics/bty560

79. Wood DE, Lu J, Langmead B. Improved metagenomic analysis with Kraken 2. Genome Biol. 2019;20(1):257. doi:10.1186/s13059-019-1891-0

80. BenLangmead. PlusPF – 16 index. Kraken 2 / Bracken Refseq indexes. Published online 2023. https://benlangmead.github.io/aws-indexes/k2

81. Valenzuela-Vallejo L, Chrysafi P, Guatibonza-García V, et al. Growth Differentiation Factor 15 (GDF-15) Levels Associated with the Presence and Severity of NAFLD. Metab - Clin Exp. 2023;142. doi:10.1016/j.metabol.2023.155445

82. Zorzano A. 2 - Intracellular Signaling Mechanisms Involved in Insulin Action. In: Serrano Ríos M, Caro JF, Carraro R, Gutiérrez Fuentes JA, eds. The Metabolic Syndrome at the Beginning of the XXI Century. Elsevier España; 2005:15–42. doi:10.1016/B978-84-8174-892-5.50002-4

